# High-resolution methylome analysis in the clonal *Populus nigra* cv. ‘Italica’ reveals environmentally sensitive hotspots and drought-responsive TE superfamilies

**DOI:** 10.1101/2022.10.18.512698

**Authors:** C. Peña-Ponton, B. Diez-Rodriguez, P. Perez-Bello, C. Becker, L.M. McIntyre, W. Van der Putten, E. De Paoli, K. Heer, L. Opgenoorth, KJF. Verhoeven

## Abstract

- DNA methylation is environment-sensitive and can mediate plant stress responses. In long-lived trees, changing environments might cumulatively shape the methylome landscape over their lifetime. However, because high-resolution methylome studies usually focus on single environments, it remains unclear to what extent the methylation responses are generic or stress-specific, and how this relates to their long-term stability.
- Here, we studied the methylome plasticity of a single poplar genotype, *Populus nigra* cv. ‘Italica’. Adult poplar trees with diverse environmental histories were clonally propagated, and the ramets exposed to experimental cold, heat, drought, herbivory, rust infection and salicylic acid treatments. Then, we identified and compared stress-induced vs. naturally occurring DNA methylation changes using whole genome bisulfite sequencing data.
- Methylation changes mainly targeted transposable elements and when occurring in CG/CHG contexts, the same regions were often affected by multiple stresses, indicating a generic response. Drought triggered a unique CHH hypermethylation response in transposable elements, affecting entire superfamilies and often occurring near drought-responsive genes. Stress-induced methylation variation in CG/CHG contexts showed striking overlap with methylation differences observed between trees from distinct geographical locations.
- Altogether, our results indicate that generic methylome stress responses can persist as epialleles in nature while some environments trigger more transient but large and specific responses, with possible functional consequences.

## INTRODUCTION

Plants are challenged by abiotic and biotic stresses that affect their survival, growth, and fitness. Long-lived trees must acclimate to simultaneous and seasonal stress exposures every year by employing diverse genetic and epigenetic strategies for regulation of plant growth, development, and reproduction. However, although the role of epigenetic mechanisms in stress responses is receiving increasing attention (Deleris et al., 2016; Lämke & Bäurle, 2017; H. Zhang et al., 2018), most knowledge comes from short-lived, annual species (Hagmann et al., 2015; Kenchanmane Raju et al., 2018; Wibowo et al., 2016). Moreover, as mitotically stable epigenetic marks have the potential to mediate plant responses to environmental changes (Becker et al., 2011; Boyko et al., 2010; López Sánchez et al., 2016; Schmitz et al., 2011), epigenetic research in perennials may be the key for understanding such roles over various time scales.

DNA methylation is the most abundantly studied epigenetic modification; it generally refers to cytosine methylation (5mC), the addition of a methyl group to the fifth position of the pyrimidine ring of a cytosine base. In plant genomes, 5mC occurs frequently in all three sequence contexts: the symmetric CpG and CHG along with the asymmetric CHH contexts (where H = A, T or C) (X. Zhang et al., 2006). Insights about the functionality of DNA methylation have been described in promoters, gene bodies and transposable elements (TEs). In promoter regions, methylation usually inhibits transcription initiation, while its function within the gene body is less clear (Bewick & Schmitz, 2017; Paszkowski & Whitham, 2001; H. Zhang et al., 2018; X. Zhang et al., 2006) but may act to quantitatively impede transcript elongation (Zilberman et al., 2007). TEs are enriched for DNA methylation and histone modifications, which are associated with transcriptional silencing (Matzke & Mosher, 2014).

*De novo* methylation in all sequence contexts is directed by small RNAs and catalysed by DOMAINS REARRANGED METHYLTRANSFERASE 2 (DRM2) in a process known as RNA-Directed DNA Methylation (RdDM) (Matzke & Mosher, 2014). Maintenance of CpG, CHG and CHH methylation are performed by METHYLTRANSFERASE 1 (MET1), CHROMOMETHYLASE 3 (CMT3)/CMT2, and DRM2/CMT2, respectively. Lack of DNA methyltransferase activity or methyl donor shortage following DNA replication result in passive DNA demethylation, while active DNA demethylation involves the activity of glycosylases which excise 5mC from all cytosine sequence contexts (H. Zhang et al., 2018).

While it is well-established that DNA methylation is responsive to environmental factors (Liu & He, 2020), its role in mediating environmental plasticity is less well understood. For instance, causality between induced methylation variation and modulation of gene expression is still a matter of debate (Bewick et al., 2019; Secco et al., 2015; Seymour & Gaut, 2020). Moreover, many plant stress responses are mediated by systemic signalling via hormones such as salicylic acid, jasmonic acid, auxin, ethylene, and abscisic acid (Karpiński et al., 2013; Wang et al., 2014). However, it is still undetermined whether different stresses could induce common methylation responses that can be explained by overlapping systemic signalling responses.

The cost of high-resolution DNA methylation analyses has been a major factor limiting the sample size within studies, leading to low statistical power for identifying DNA methylation variants. More importantly, this has also led to the study of single environmental factors at a time, restricting the comparison of identified responsive loci among experiments. Hence, the present view is that DNA methylation responses can be highly specific with respect to environmental conditions, which might underestimate the possibility of overlapping responses that generally occur under a variety of stresses.

To study molecular mechanisms of trees in response to environmental cues, *Populus* species have become a choice model system due to their rapid growth, easy propagation, and available genomic resources (Jansson & Douglas, 2007; Tuskan et al., 2006). Moreover, poplars are riparian species that are among the woody plants most sensitive to water stress (Larchevêque et al., 2011; Rood et al., 2003). This prompted much research on molecular mechanisms of drought tolerance (Jia et al., 2016; Viger et al., 2016; Yıldırım & Kaya, 2017), including DNA methylation (Lafon-Placette et al., 2018; Sow et al., 2021). *Populus. nigra* cv. ‘Italica’, also known as the Lombardy poplar, is one of the most widely distributed poplar cultivars that was first reported in Lombardy, Italy, at the very beginning of the eighteenth century (Chenault et al., 2011). Afterwards, it was introduced into France, from where it is believed that Napoleon promoted its spread across the Empire by clonal propagation (Stettler, 2009). As a result, Europe has become colonized by a genetically homogeneous male clone of the Lombardy poplar. The clonality of this cultivar makes it an excellent system for studying epigenetic plasticity in response to different environmental conditions, as it strongly limits the confounding effects of genetic variability (Díez-Rodríguez et al., 2022).

Here, we used whole genome bisulfite sequencing to capture the methylation responses of young poplars, clonally propagated from adult trees from different European locations, to a variety of biotic and abiotic stresses. With this unique approach, we aimed to characterize the environmentally induced methylome variation of the Lombardy poplar and to examine methylation variation as induced by exposure to acute stress in comparison with methylation variation that had built up naturally between trees during the lifetime of growth in different geographic locations. Our study revealed commonalities between the methylation responses to different stresses, very specific responses to some stresses, and hints at the long-term stability of such responses.

## MATERIALS AND METHODS

Our experimental approach involved whole genome evaluation and multiple treatments. Here, we provide a brief overview of the employed methods while a detailed methodology for each section can be found as supporting information.

### Plant material

Cuttings from eight adult *Populus nigra* cv. ‘Italica’ clones were collected from five European countries (Table S1, Figure 1A; see (Díez-Rodríguez et al., 2022). At each site, at least seven hardwood cuttings of approximately 30 cm length were sampled from each adult parental tree (ortet) and stored at 4 °C for two weeks prior to planting. Cuttings were grown under controlled conditions for 12 weeks until the start of the experiment. Growth conditions were: 22/18 °C (±2°C) at day/night, 60% relative humidity (±5% Rh), 16/8 h light/dark. Cuttings were planted first in 4-liter pots with a 3:1 sand:peat mixture (v/v) and placed in a flood table for three weeks. Rooted cuttings (ramets) of similar size were transferred to 7-liter pots with a 1:1 sand:peat mixture and maintained with regular watering. Two weeks prior to the start of the experiment, three grams of slow-release fertilizer Osmocote Exact Mini (16+8+11+2MgO+TE) were added to each pot.

**Figure 1.**
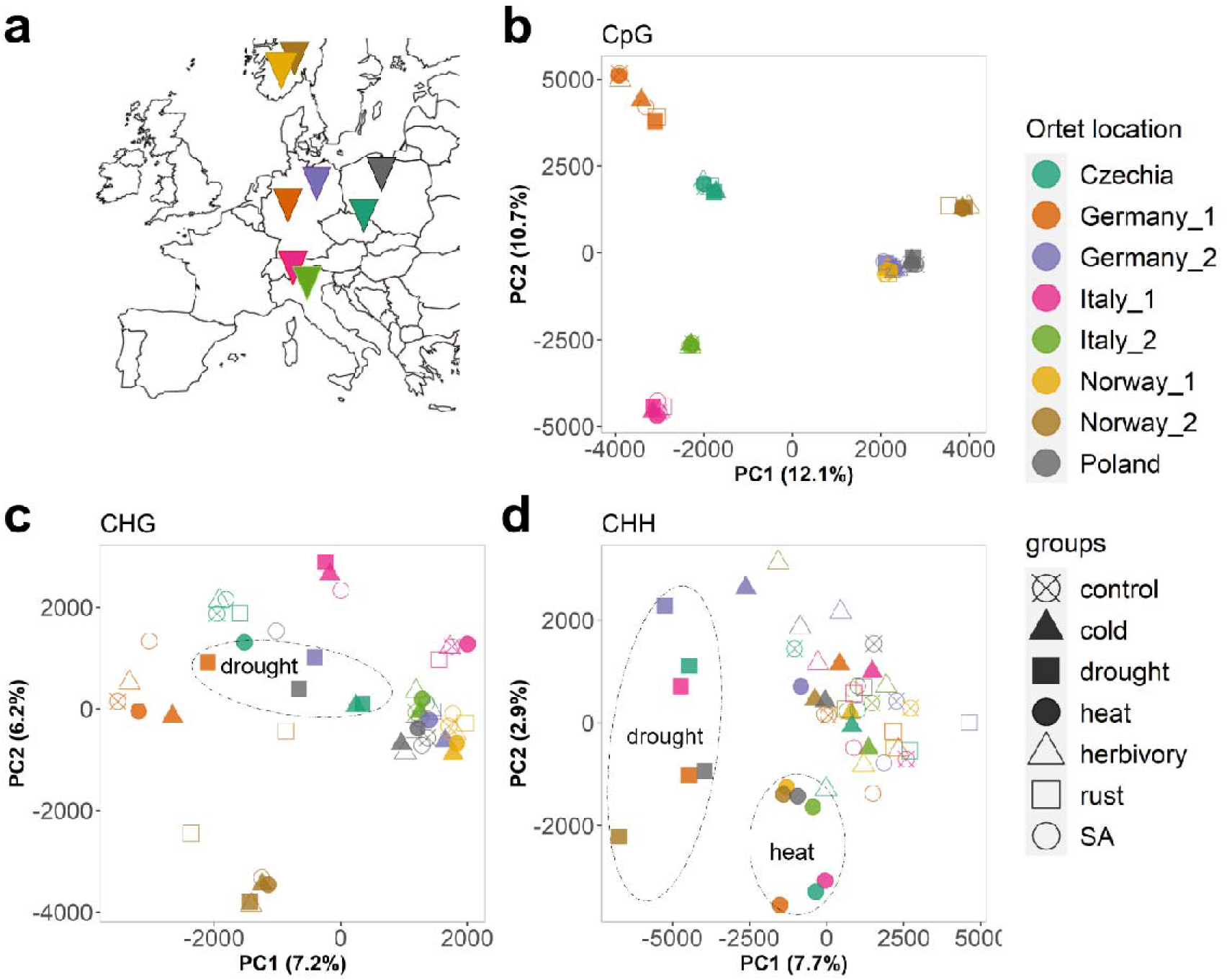
Analysis of genome-wide methylation patterns from stress-treated Lombardy poplar ramets. **a)** Sampling locations of the clonally propagated ortets used for the experiment. **b**, **c**, and **d)** Unsupervised principal component analysis of CpG, CHG and CHH methylation. Ramets are coloured by ortet identity. Different shapes represent each experimental group. Drought and heat clusters are highlighted with dashed ovals.

### Experimental design

Seven 3-month-old ramets of similar size were selected from each of the eight (ortets) for subsequent exposure to different environmental treatments (56 trees in total). The experiment consisted of three biotic and three abiotic stresses plus a single control group, with eight replicates per treatment, where each ortet contributed one replicate to each of the treatments. All treatments were implemented simultaneously during a period of 25 days, including two 10-day stress events and one 5-day stress-free period in between (Figure S1).

#### Control group

during the entire stress experiment, control plants were maintained in a greenhouse under controlled conditions as stated above. The soil volumetric water content (VWC) was maintained on average at 20.2% (± 3.24 SD) by daily watering to pot capacity. VWC was monitored daily using the WET Sensor kit (Delta-T Devices). Mean VWC was calculated for all plants with two measurements per pot (Figure S2). During the entire experiment, control plants were maintained in the same greenhouse table with other stress treatments in a Latin square design unless otherwise stated (Figure S2).

#### Biotic stresses

Rust infection consisted in spray-inoculation of uredospores of the poplar leaf rust fungus (*Melampsora larici-populina* Kleb.). Herbivory treatment involved the use of Gypsy moth caterpillars (*Lymantria dispar* L.). Salicylic acid (SA) treatment consisted in spray application of 1 mM SA (Sigma-Aldrich). Rust spores and caterpillars were obtained from Dr. Sybille Unsicker (Max Planck Institute for Chemical Ecology, Jena, Germany).

#### Abiotic stresses

Drought stress was attained by withholding watering and maintaining VWC at 8%. Plants that received heat treatment were grown at high temperatures of 30-38/28°C (day/night), while cold-treated plants were grown at 4/4°C (day/night).

VWC for all treatments (except drought) was maintained close to control conditions (Fig. S3a). Plants were moved to climate chambers for heat and cold treatments (Fig. S3b). During the stress-free period, plants were moved to the same greenhouse table as the control group (Fig. S2).

#### Harvesting

For DNA methylation analysis, on experimental day 26, twelve circular punches (Ø 8 mm; ~ 100 mg fresh weight in total) were cut out from the eighth mature leaf (counting from the apex of the main branch, leaf plastochron index: 10) of each plant. MidLribs were avoided, and leaf punches were immediately frozen in liquid nitrogen and stored at −80 °C. All sampled leaves were not directly exposed to any of the biotic stresses; thus, we characterized the systemic response.

### DNA extraction and whole genome bisulfite sequencing

Per sample, frozen leaf tissue was grinded and homogenized using TissueLyser II (QIAGEN), then genomic DNA was isolated using the Sodium Dodecyl Sulfate (SDS) procedure of the NucleoSpin Plant II DNA isolation kit (Macherey-Nagel, Dueren, Germany). Preparation of DNA libraries for bisulfite sequencing was performed as described in (Nunn et al., 2022). All sequencing was performed by Novogene on an Illumina HiSeq X Ten sequencing system. Libraries were sequenced with 2×150-bp paired-end reads at 30X coverage. Libraries were sequenced in a total of eight sequencing lanes, trying to allocate ramets derived from the same ortet in the same lane to avoid batch effects.

### Processing of bisulfite-treated reads and methylation calling

Sequenced reads were processed using the EpiDiverse Toolkit (WGBS pipeline v1.0, https://github.com/EpiDiverse/wgbs) (Nunn et al., 2021). Briefly, low-quality read-ends were trimmed (minimum base quality: 20), sequencing adapters were removed (minimum overlap: 3 bp) and very short reads (< 36 bp) were discarded. The remaining high-quality reads were aligned to the *Populus nig*r*a* var. ‘Italica’ *de novo* reference genome (ENA project: PRJEB44889) using erne-bs5 (http://erne.sourceforge.net) allowing for 600-bp maximum insert size, 0.05 mismatches, and unique mapping. Per-cytosine methylation metrics were calculated using MethylDackel (https://github.com/dpryan79/MethylDackel). Three bedGraph files per sample were obtained, corresponding to cytosines on each sequence context: CpG, CHG and CHH. The methylation level (%) of a particular site was calculated by:

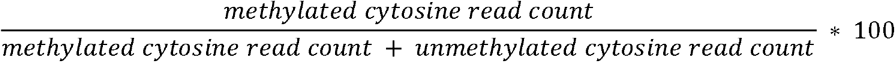

### Methylation analysis

The analysis followed three stages: 1) genome-wide methylation analysis: to detect strong global methylation patterns by comparing samples and groups to each other, 2) differential methylation analysis: to identify significant differentially methylated regions (DMRs) by testing for significant methylation differences between groups throughout the entire genome, and 3) downstream analysis: to reveal potential functional implications by testing annotated DMRs for enrichment on genomic features and gene functions. The three cytosine sequence contexts were always analysed separately. Data filtering and resolution was slightly different for each analysis (Table S2). As a first general filtering step, all cytosines with low sequencing coverage (≤ 5 reads) were removed.

### Genome-wide methylation analysis

After the first filtering step, samples in which the retained cytosines accounted for less than 50% of the original data were considered low-coverage outliers and were excluded from genome-wide analyses unless otherwise stated (4 outliers out of 56 samples, Supplementary file 1, Table S2).

### Average global methylation

Only genomic positions with methylation information across all 52 remaining samples were considered (CpG: 1’802.288 positions, CHG: 3’256.938 positions, CHH: 12’058.984 positions). For each sample, average global methylation (%) was calculated separately for each context, as follows:

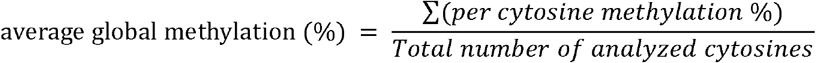

The effect of the stress treatments on the level of global methylation was evaluated for each context using a linear mixed model with treatment as fixed factor and parental tree (ortet) as random factor. Statistical analyses were calculated in R (version 4.0.3), the *lmer* function of the *lmerTest* package (Kuznetsova et al., 2017) was used to fit the model, and multiple comparisons (Tukey’s post-hoc tests) were calculated with the *glht* function of the *multcomp* package (Hothorn et al., 2008).

### Principal component analysis, hierarchical clustering, and correlation analysis

For CpG and CHG context, the same filtered data used for average global methylation analysis was considered for principal component analysis (PCA) and hierarchical clustering (HC). As we observed that a very large fraction of cytosines in CHH context showed very low methylation variation across samples, an extra filtering step removed CHH positions where more than 90% of the samples showed very low (0-5%) or very high (95-100%) methylation to keep the more variable positions.

Principal components (PCs) were calculated in R using the *prcomp* function of the *stats* package (R Core Team, 2022). HC (Ward’s method) was computed by first calculating the corresponding distance matrix (Manhattan method) using the *dist* and *hclust* functions from the *stats* R package.

Pairwise correlation analysis was performed using genomic regions instead of single positions. First, the poplar genome was compartmentalized in 100-bp non-overlapping bins. Average methylation per bin was calculated, and only bins with methylation information across all samples were retained. As the intraclass correlation coefficient (ICC) reflects both degree of correlation and agreement between measurements (Koo & Li, 2016), ICC was calculated for all pairwise comparisons between samples. Coefficients were calculated in R using the *icc* function of the *irr* package (Gamer et al., 2019) with the ICC form: two-way random effects, absolute agreement, single measurement, according to (McGraw & Wong, 1996) convention.

### Methylation profiles

All (56) samples were included in the analysis. For gene regions, only protein-coding genes with known 5’UTR and 3’UTR coordinates were considered. For transposable elements, only TEs longer than 150 bp were analyzed. Methylation profiles over the largest poplar scaffold (scaffold 1 = 33’746.648 bases) were used as a proxy for chromosome-wise methylation variation comparison. The scaffold was compartmentalized in 50-kb bins, then for each sample, per-bin average methylation was calculated. Finally, to calculate per-bin methylation for each treatment, a weighted mean was calculated accounting for the number of cytosines per bin per sample using the formula:

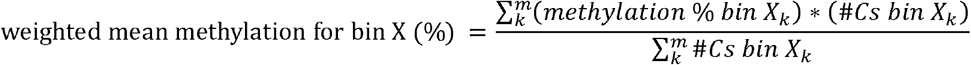

where ***X*** represents any 50-kb bin, ***k*** is a replicate, and ***m*** is the total number of replicates per treatment

For each treatment and context, per-bin methylation differences compared to the control group were calculated and used to plot heatmaps and simple moving averages (SMA). SMAs were calculated using the R function *geom_ma* of the *tidyquant* package (Dancho, 2022).

### Differential methylation analysis

Differentially methylated regions (DMRs) induced by each stress treatment were identified by testing local methylation differences between each treatment and control group. All replicates per treatment were included in the tests and each cytosine sequence context was analysed separately using the EpiDiverse/DMR pipeline v0.9.1 (https://github.com/EpiDiverse/dmr) (Nunn et al., 2021). Briefly, DMRs were identified by metilene (https://www.bioinf.uni-leipzig.de/Software/metilene), with parameters as follows. Minimum read depth per position: 6; minimum cytosine number per DMR: 10; minimum distance between two different DMRs: 146 bp; per-group minimal non-missing data for estimating missing values: 0.8; adjusted p-value (Benjamini-Hochberg) to detect significant DMRs: 0.05. Only significant DMRs with minimum methylation difference of 10 percentage points between groups were used for downstream analyses.

### DMR calling

Since the genome-wide methylation analyses revealed strong CpG and CHG methylation patterns associated to sample origin (ortet identity) irrespective of stress treatment, stress-DMRs were identified using a jack-knife approach (leave-one-out) to reduce within-treatment variation caused by individual outlying ortets. As eight different ortets were included in the experimental design, a total of eight DMR calls were performed, in which samples derived from single ortets were left out on each DMR call (Figure S4). Hence, seven replicates were included on each DMR calling. All identified DMRs were retained for further analysis. Additionally, to check if our jack-knife approach for DMR calling produced robust results, we called stress-DMRs using Methylkit (Akalin et al., 2012) and intersected both results. This check indicated a large overlap between both methods and highlighted the conservative nature of our results (Table S3).

Based on overlaps among DMRs that were identified in more than one treatment, DMRs were classified as multi-stress or stress-specific DMRs. Genomic regions where a DMR was identified in more than one sequence context were labelled as multi-context DMRs.

We also performed DMR callings among ortets. Ramets derived from the same ortet and exposed to different treatments were considered replicates. Briefly, DMRs were called for all pairwise comparisons among the eight ortets (total: 28 DMR-sets per context). Next, for each context, all DMRs were classified according to the number of pairwise comparisons in which each DMR appeared in: DMRs found in a *unique comparison* or DMRs *shared by two or more comparisons*.

### Downstream methylation analyses

#### DMR annotation

Statistically significant DMRs were annotated using the *Populus nigra* cv. ‘Italica’ protein-coding gene model annotation. Only the longest transcript per gene was used for this analysis. DMRs were also associated with TEs based on a TE prediction for this cultivar. Gene models and TE predictions used in this study were generated as part of the ongoing *P. nigra* cv. ‘Italica’ reference genome project (PRJEB44889). Short descriptions of these annotation files can be found along with their deposited versions (see Data availability statement). Short interspersed nuclear elements (SINEs) were manually added to the predicted TEs based on BLASTN results (70% similarity, 90% coverage) using the consensus sequences of Salicaceae SINE families (Kögler et al., 2020).

#### DMR enrichment on genomic features

For each context, all DMRs, irrespective of treatment, were tested for enrichment in gene bodies, exons, introns, gene flanking regions and TEs (Z-test for proportions). As DMRs were enriched in TEs, we also tested whether the occurrence of DMRs in gene bodies, exons, introns and gene flanking regions was conditional on the presence of TEs (Chi-square tests for independence, McNemar’s test). Additionally, as drought showed the largest response associated to TEs, we calculated the relative fold enrichment for drought CHH-DMRs on each TE superfamily. P-values were obtained from the hypergeometric test and then adjusted (Bonferroni) according to the number of TE superfamilies tested. Drought-DMR-enriched TE superfamilies were referred to as Drought-Responsive TEs (DR-TEs), which included all SINE and MITE/DTHs elements.

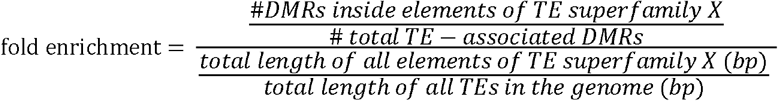

#### Stress-induced methylation variation of DMRs and TEs

Since stress-DMRs were identified between each treatment vs. the control group, we were interested in evaluating the methylation response of such regions induced by all the other treatments. Using the genomic coordinates of the identified DMRs, we calculated average methylation levels of the corresponding regions on each sample Then, we calculated the average methylation level across treatment replicates. Finally, for each region, we computed the methylation difference between each treatment and control group. We only analysed regions with enough methylation information (≥ 8 cytosines) and replication (≥ 6 replicates).

Moreover, as we detected TE superfamilies enriched with drought-DMRs, we analysed the drought-induced methylation response of all poplar TE superfamilies to check for generalized responses of entire TE superfamilies. For each sequence context, average methylation levels of each individual TE were calculated for drought and control samples. We only analysed TEs with enough methylation information (≥20 cytosines for CHH, ≥10 cytosines for CpG and CHG) and replication (≥ 6 replicates). For each TE element, we computed the methylation difference between drought and control group. Then, to summarize and compare results among TE superfamilies, we grouped TE elements in boxplots according to each superfamily.

#### Gene ontology enrichment analysis

Functional enrichment analysis has to be carefully interpreted as gene expression data was not collected in this experiment and most stresses produced very few DMRs. Therefore, our analysis was mainly focused on medium-to-large gene sets associated with drought-induced methylation responses.

Genes associated with drought CHH-DMRs were subjected to gene ontology (GO) enrichment analysis. The gene background was built with the closest Arabidopsis (*A. thaliana*) homologue of each *P. nigra* cv. ‘Italica’ gene, which was determined using BLAST best reciprocal hits (RBH) of the protein sequences (R package *orthologr* (Drost et al., 2015). Best hits were filtered by keeping alignments covering at least 60% of both *Arabidopsis* and *P. nigra* proteins, and minimum 60% similarity. *Arabidopsis* protein sequences were extracted from phytozome V13, and functional annotations were retrieved from the PLAZA 5.0 dicots database (https://bioinformatics.psb.ugent.be/plaza/). GO enrichments were performed using clusterProfiler v4 (Wu et al., 2021). P-values were adjusted for multiple testing controlling the positive false discovery rate (q-value).

Enrichments for genes associated with all SINE and MITE/DTH elements were performed in the same manner. Gene sets for functional enrichment included genes associated with either all SINEs, all MITE/DTHs, or both (DR-TEs). Additional enrichments were performed for a subset of potential Highly Drought-Responsive TEs (HDR-TEs), i.e., SINEs and MITE/DTHs that displayed at least 5% hypermethylation compared to the control group. Finally, for comparison, all enrichments were analysed in the context of the drought CHH-DMR gene set enrichment.

## RESULTS

### Drought induces a large and distinctive genome-wide CHH hypermethylation response in the Lombardy poplar

Genome-wide methylation analyses were performed to identify overall strong methylation patterns among samples and treatments. Starting with average global methylation, the linear mixed models revealed significant treatment effects on DNA methylation (CpG: p=0.048, CHG: p=0.043, CHH: p<0.01). However, only drought treatment induced a significant global increase of CHH methylation compared to control group (Tukey’s test p<0.01). In addition, cold treatment induced significantly higher CpG and CHG methylation levels compared to salicylic acid treatment (p=0.0241 and 0.0412, respectively) (Fig. S5a).

For CHH methylation, PCA and HC analyses highlighted noticeable clusters for drought and heat treatments (Fig. 1d, S6c, S7b). High correlations were observed among drought-treated samples, while the lowest correlation coefficients were found when comparing drought samples with any other sample (Figure S8c). Methylation profiles over the largest poplar scaffold confirmed the genome-wide drought-induced CHH hypermethylation and underlined a close relationship with TE content as both profiles showed peak similarities (Fig. 2, S9). Over TE regions, profiles corroborated the drought-induced CHH hypermethylation, and highlighted CHH hypomethylation induced by rust infection. Profiles over genic regions revealed that drought-induced CHH hypermethylation mainly targeted gene-flanking regions rather than gene bodies (Fig S10, S11).

**Figure 2.**
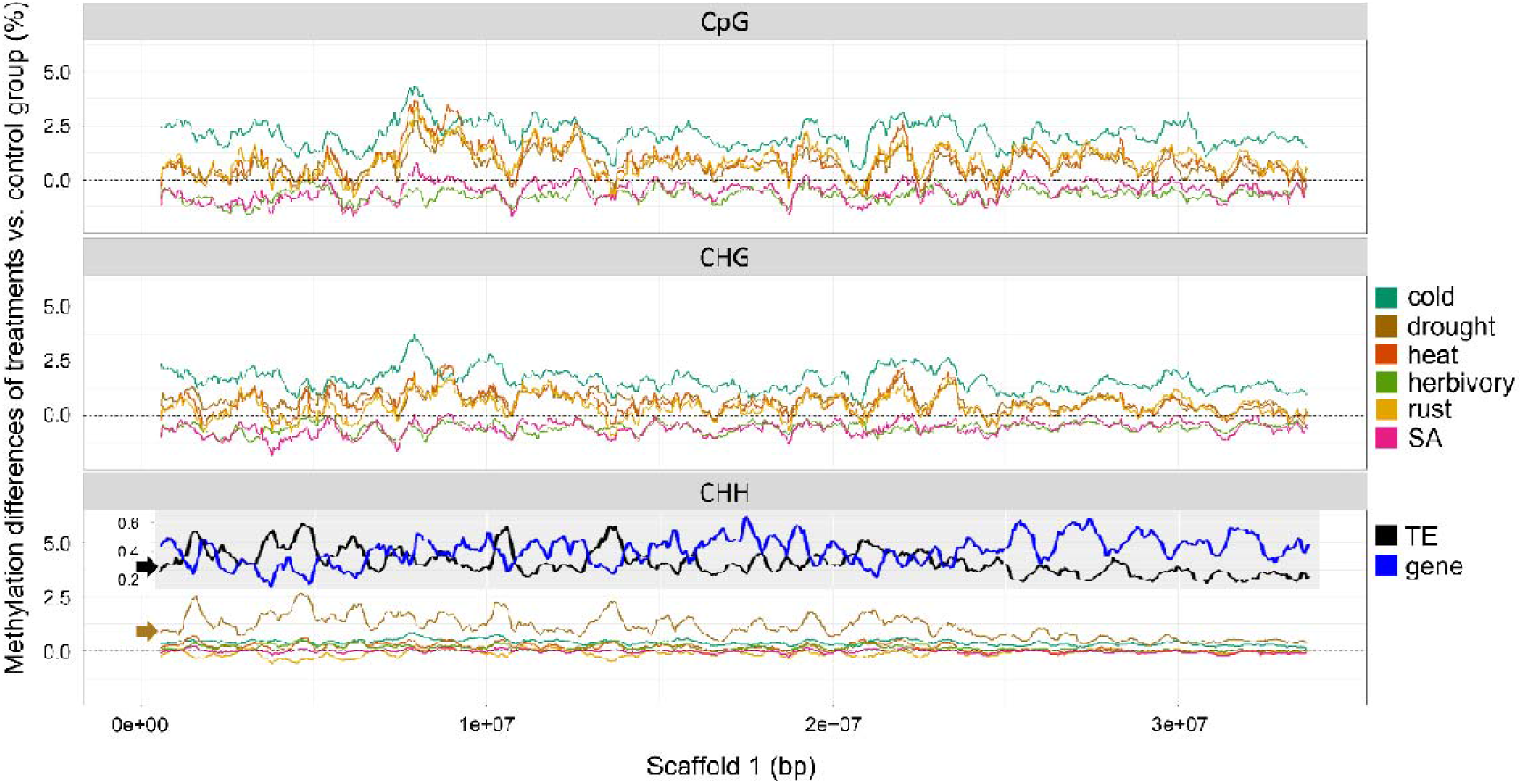
Metaplots of CpG, CHG and CHH methylation level differences (treatments vs control group) over the scaffold 1 of *Populus nigra* var. Italica. Simple moving averages (SMA) over a period of ten 50-kb bins were calculated and plotted for each treatment and context. Profiles for TE and gene content were added on top of CHH metaplot (SMAs per ten 50-kb bin) to highlight the relationship between methylation variation and gene/TE content, especially obvious for drought-induced CHH methylation variation (brown arrow) and TE content profile (black arrow).

The effect of other treatments was also observed in the methylation profiles. Profiles of CpG/CHG methylation along the scaffold 1 confirmed the genome-wide cold-induced hypermethylation that was already detected in the global methylation analysis. In addition, several treatments showed overlapping profiles of hypermethylation (drought, heat, and rust), and hypomethylation (SA and herbivory). Visual observation of the methylation profiles indicated a positive correlation between CpG and CHG methylation variation (Fig. 2, S9). Interestingly, over genic regions, the effects of stress treatments were detectable mainly in gene flanking regions while in TE regions, cold and SA induced the largest CpG/CHG methylation responses: hypermethylation and hypomethylation, respectively (Figure S11).

### Genome-wide CpG and CHG methylation largely reflect sample origin rather than treatment effect

Linear mixed models revealed significant ortet effects on the average global CpG methylation (p<0.01) (Figure S5b). Detailed insights were observed on PCA, HC and correlation analysis where distances among ramets derived from the same ortet (within-ortet) were much smaller than those among ramets derived from different ortets (between-ortet), irrespective of the treatment (Figure 1b, S6a, S8a, S8d). A similar but less pronounced within-ortet clustering was found for CHG methylation, with an additional clustering of droughtlJtreated ramets, irrespective of the ortet (Figure 1c, S6b, S7a, S8e).

### Transposable elements are enriched with stress-induced DMRs

We identified a total of 1,798 DMRs across all treatments and sequence contexts (Fig. 3, table S4). Drought induced the largest number of DMRs among all treatments (mostly CHH hypermethylations), while cold induced the largest amount of hypermethylated DMRs in CpG/CHG contexts (Fig. 3a, table S4). In general, similar amounts of multi-stress and stress-specific DMRs were observed in each treatment, except for drought CHH-DMRs (Figure 3b, Table S5). Among the 203 multi-stress DMRs, drought and heat showed the largest intersection with 57 DMRs (Fig. S12).

**Figure 3.**
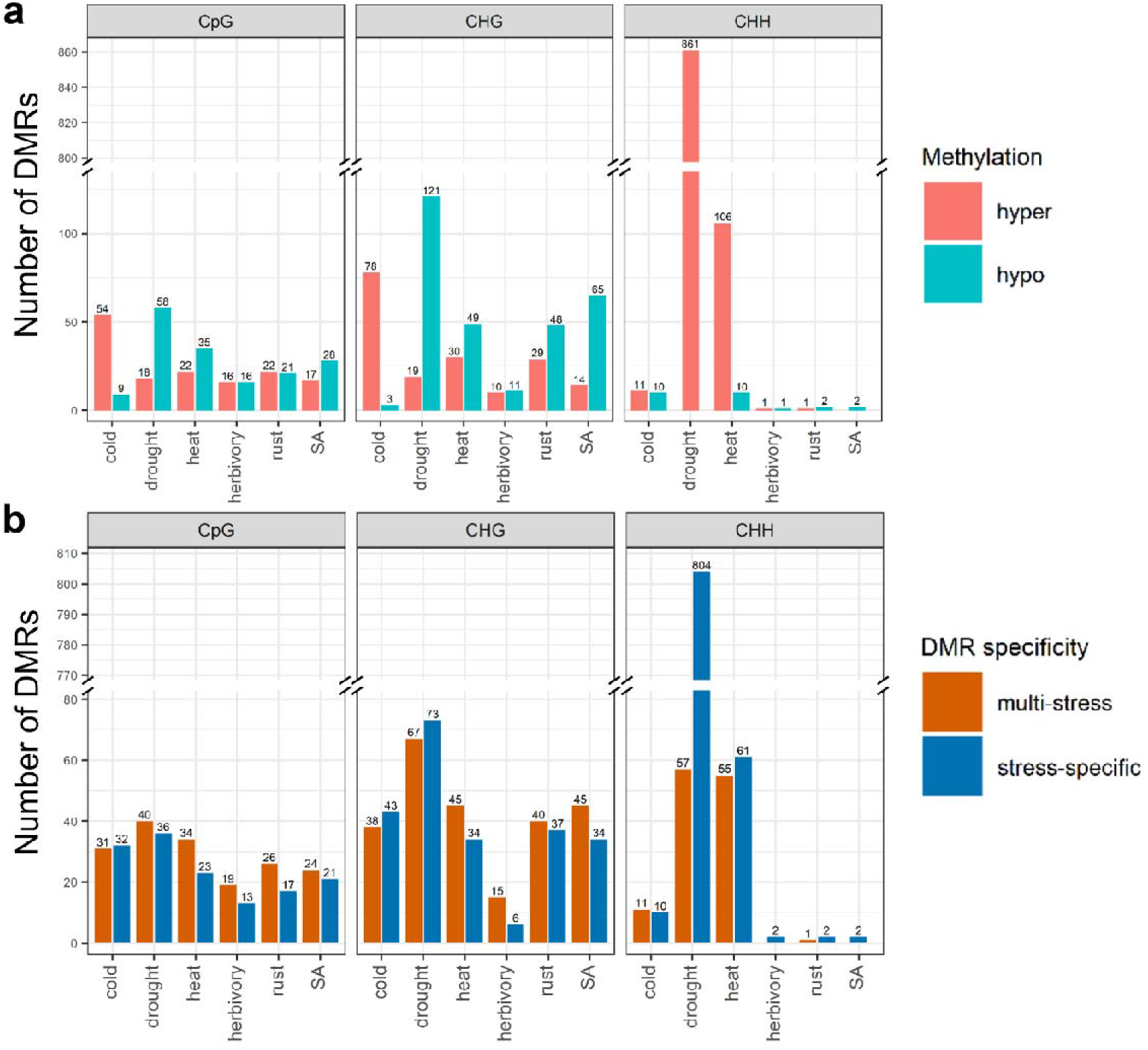
Summary of significant DMRs induced in the Lombardy poplar by each stress treatment versus the control group in CpG, CHG and CHH contexts. **a)** DMRs classified by methylation direction: hypermethylated (red), hypomethylated (blue) **b)** DMRs classified by specificity: multi-stress (orange), stress-specific (dark blue). DMR numbers are shown on top of each bar.

Enrichment tests showed that all DMRs, irrespective of sequence context, were enriched in TEs. In addition, CpG-DMRs mainly targeted gene bodies, specifically exons, while CHH-DMRs were enriched over gene flanking regions (Fig. 4b, Table S6). CHG-DMRs were enriched in intergenic regions associated with TEs, while TE-associated CHH-DMRs were enriched in gene flanking regions and introns (Table S7). Moreover, we observed an increased frequency of DMRs in TE flanking regions, especially within the first 200 bp (Fig. S13), revealing TEs as a major source of methylation variation (DMRs) irrespective of treatment (Fig. S18, S19, S20)

**Figure 4.**
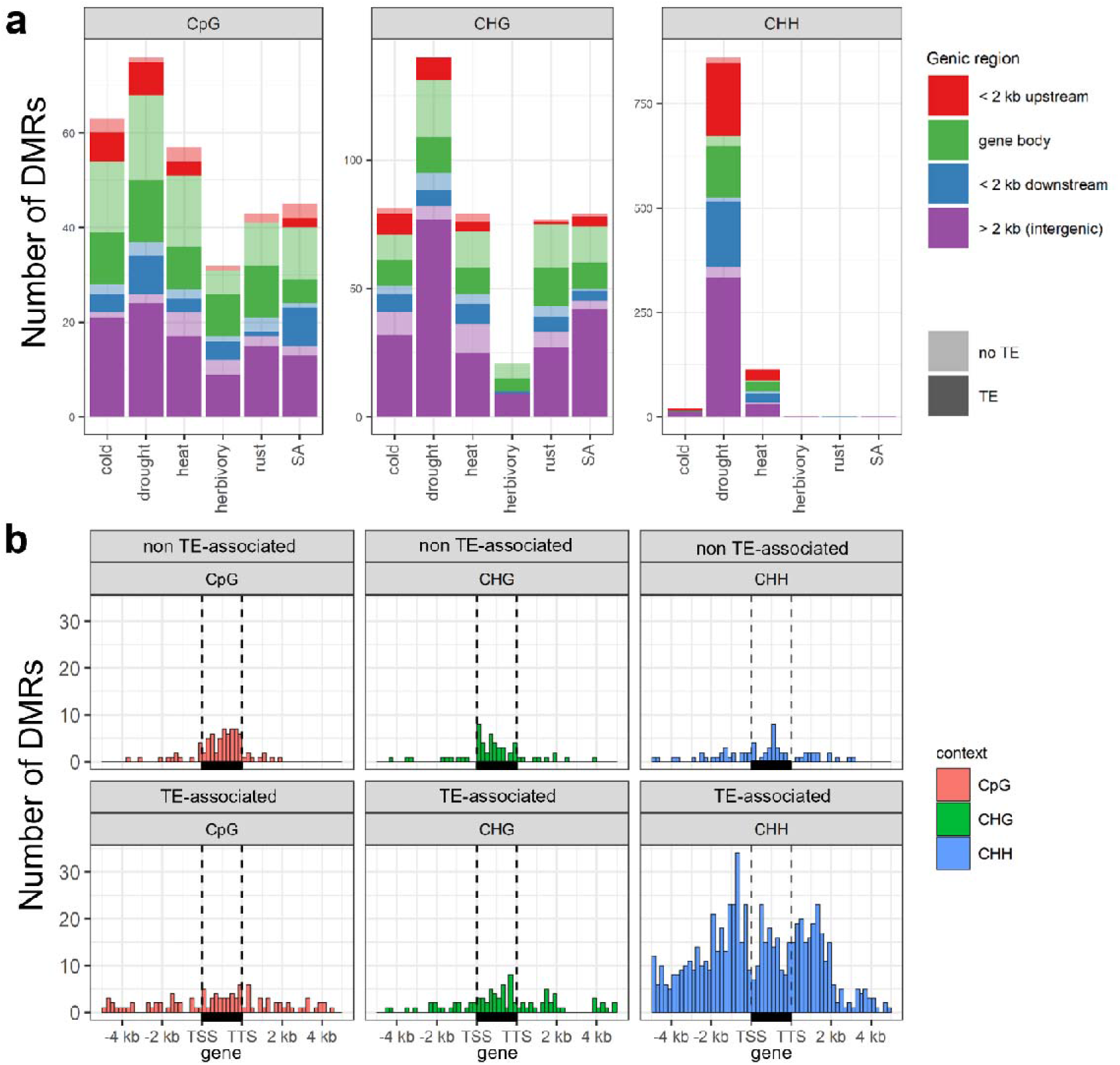
Distribution of stress-induced DMRs over the Lombardy poplar genome. **a)** For each treatment and context, DMR counts (irrespective of treatment) are shown for gene body, ±2kb gene flanking regions, and intergenic regions. Dark/light colors differentiate the number of DMRs associated/non-associated with TEs in the corresponding region. **b)** Detailed distribution of all stress-induced DMRs along genic regions, per context and TE association. Vertical dashed lines indicate the gene transcription start site (TSS) and transcription termination site (TTS). Horizontal black boxes represent the gene body. Gene lengths were normalized to 2kb. Z-tests for proportions were performed based on the content of each genomic feature in the poplar genome (See Table S6 for complete results).

### Drought-induced CHH hypermethylation is stronger on specific TE superfamilies

DMR enrichments over each TE superfamily revealed that Short Interspersed Nuclear Elements (SINE) and Miniature Inverted-repeat Transposable Elements (MITE), especially MITE/DTH, showed an exceptionally strong response (Fig. 5, Table S8). Interestingly, SINE and MITE/DTH elements also display the highest CHH methylation under control conditions among all TE superfamilies (Figure S14). Methylation analysis over genic regions showed that drought induced CHH hypermethylation of SINE and MITE/DTH elements irrespective of gene proximity (Fig. S15 and S16, respectively).

**Figure 5.**
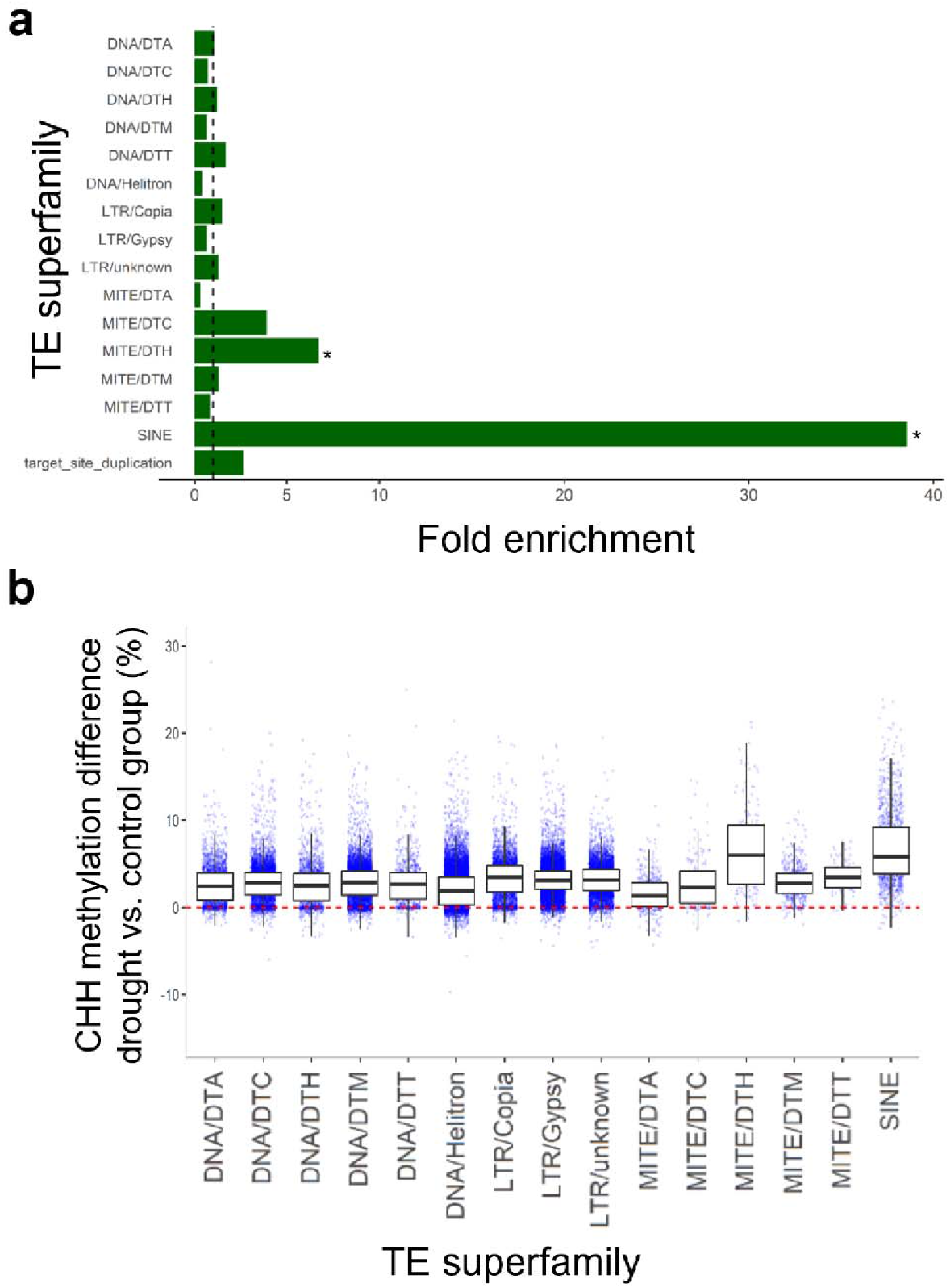
Analysis of drought-induced CHH hypermethylation of the Lombardy poplar TE superfamilies. **a)** Fold enrichment analysis of TE superfamilies targeted by drought CHH-DMRs. Enrichments were calculated based on the total length of each TE superfamily in the genome. Hypergeometric tests identified significant enrichments (p<0.001) for SINE and MITE/DTH superfamilies (Table S10). **b)** Boxplots of drought-induced CHH methylation variation (vs. control group) over individual TE elements. Each boxplot summarizes the overall methylation response of a specific TE superfamily (x-axis) to drought stress. Horizontal dotted red line depicts the zero drought-control difference.

### CpG/CHG stress-DMRs are also multi-stress DMRs and ortet-DMRs

By examining the methylation response of stress-specific DMR regions in all other treatments, we observed that most of the stress-specific CpG/CHG-DMRs also showed a response to other treatments, usually in the same direction (either hyper- or hypo-methylation) (Fig. 6a, S17). Thus, different stresses tended to result in similar methylation responses at these genomic locations, even when statistical significance was only reached in response to some treatments. Moreover, by examining the methylation level of these responsive regions in the control group, we noticed that CpG/CHG-DMRs had intermediate CpG/CHG methylation and low CHH methylation, while CHH-DMRs showed very high methylation in all contexts (Fig. 6b).

**Figure 6.**
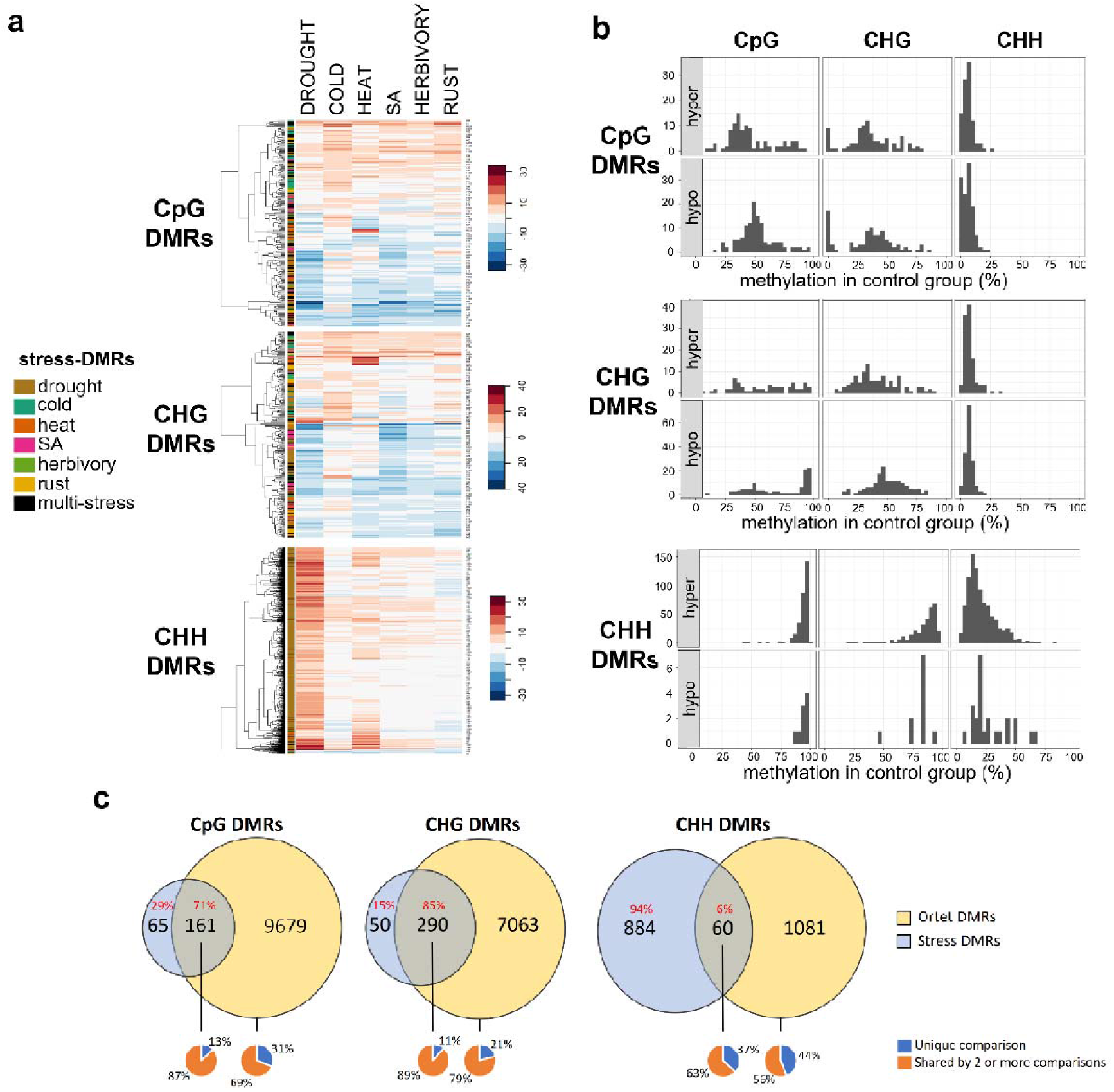
Detailed methylation patterns of DMRs identified in the Lombardy poplar. **a)** Heatmap and hierarchical clustering of the average difference methylation levels (compared to control) of the 1,728 identified stress-DMRs **b)** Histograms of CpG, CHG and CHH methylation level in the control group for all stress-DMRs. Histograms are shown according to DMR features (context and response: hyper/hypo). **c)** Venn diagrams of the intersections between ortet-DMRs and stress-DMRs for each sequence context. Uniqueness of ortet-DMRs (and for the intersection) is shown below the Venn diagrams.

The amount of ortet-DMRs was several orders higher than the stress-DMRs, especially in CpG/CHG context. For each comparison between two ortets, we identified on average 1,425 CpG-DMRs, 1,621 CHG-DMRs and 133 CHH-DMRs (Table S9). We detected a total of 9,840 CpG-DMRs, 7,353 CHG-DMRs and 1,141 CHH-DMRs after accounting for DMRs found in more than one pairwise comparison (Table S10). Such ortet-DMRs were considered as a product of natural methylation variation, and when intersected with stress-DMRs, the analysis revealed that most of the stress CpG-DMRs (71%) and CHG-DMRs (85%) were also identified as ortet-DMRs. However, only 6% of stress CHH-DMRs were found in the ortet-DMR set. (Fig. 6c, Table S10).

### Functional analysis of genes associated to drought-DMRs and drought-responsive TEs

Enrichment analyses revealed very few gene ontology (GO) terms that were significant after multiple testing correction (q-value). GO terms with uncorrected p-values (<0.05) suggested that genes associated with drought CHH-DMRs were enriched in processes related to response to abiotic stimulus (GO:0071214), osmotic stress (GO:0006970), and water deprivation (GO:0009414) (Supplementary file 2). Comparisons of functional enrichments of gene sets associated to drought CHH-DMRs and drought-responsive TEs (DR-TEs) highlighted considerable overlaps. Response to abscisic acid (GO:0009737) and protein kinase activity (GO:0004672) were terms that were enriched in almost all gene sets, while cellular response to water deprivation (GO:0042631) and cellular response to water stimulus (GO:0071462) were enriched only in drought-DMR, MITE/DTH and DR-TE sets. In addition, SINE-associated genes were mainly enriched in terms related to protein phosphorylation while MITE/DTH-associated genes were mostly enriched in terms related to ABA/hormone signalling and response to water stimulus. Gene sets associated with HDR-TEs showed similar enrichments than those accounting for DR-TEs (Table 1). Thus, regions and TE superfamilies that showed methylation responses to drought seem to be located close to drought-responsive genes.

**Table 1.**
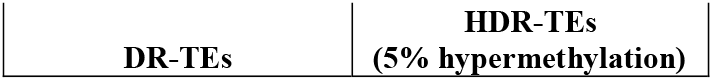
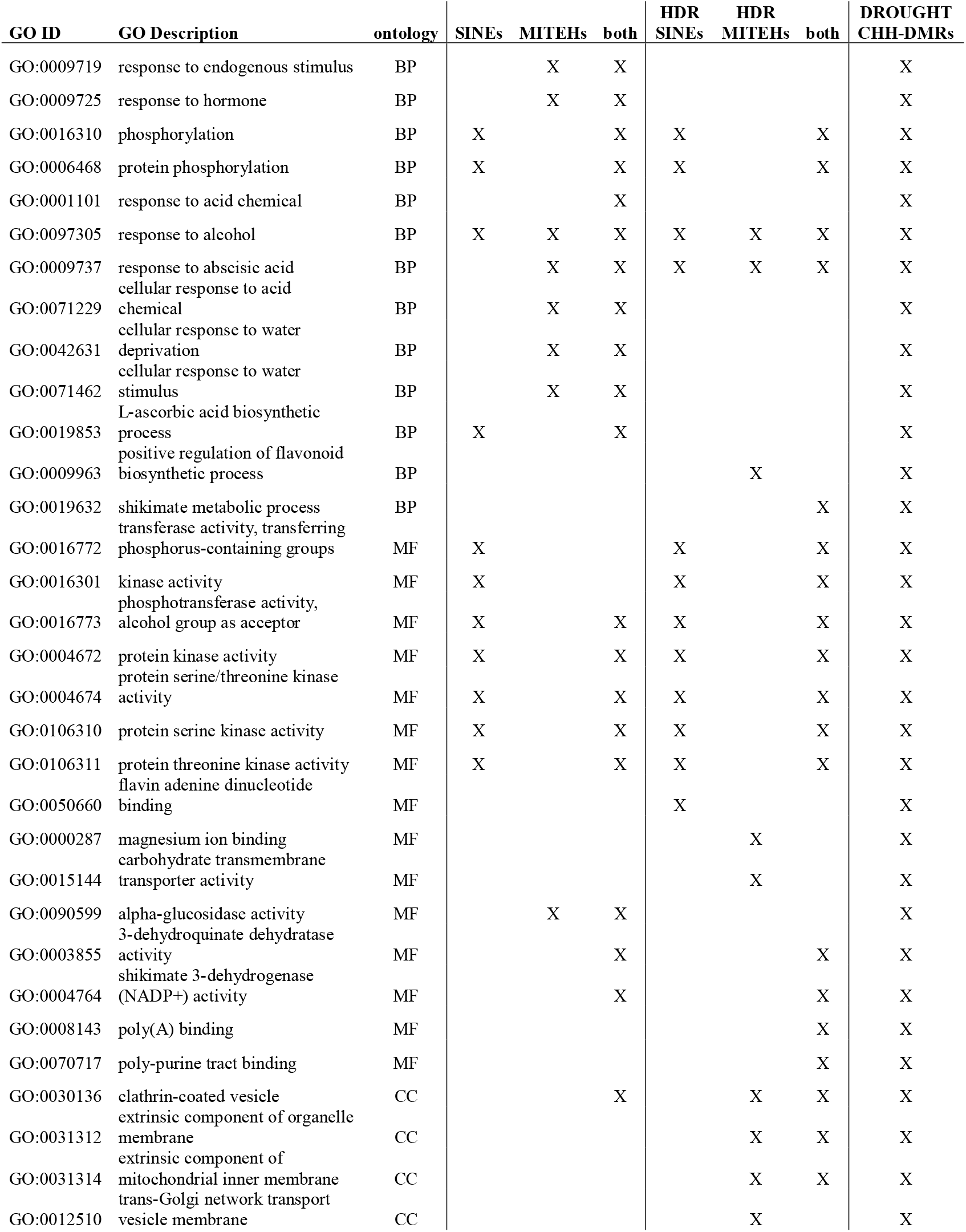
Gene Ontology (GO) enrichment analysis of genes associated with drought CHH-DMRs and Drought-Responsive TEs (DR-TEs) and Highly Drought-Responsive TEs (HDR-TEs). Only significant GO terms (p-value < 0.05) for each gene set are marked by “X”. Enriched GO terms for drought CHH-DMRs (far right) were used as a basis for comparison among all gene set enrichments.

| GO ID | GO Description | ontology | SINEs | MITEHs | both | HDR<br>SINEs | HDR<br>MITEHs | both | DROUGHT<br>CHH-DMRs |
| --- | --- | --- | --- | --- | --- | --- | --- | --- | --- |
| GO:0009719 | response to endogenous stimulus | BP |  | X | X |  |  |  | X |
| GO:0009725 | response to hormone | BP |  | X | X |  |  |  | X |
| GO:0016310 | phosphorylation | BP | X |  | X | X |  | X | X |
| GO:0006468 | protein phosphorylation | BP | X |  | X | X |  | X | X |
| GO:0001101 | response to acid chemical | BP |  |  | X |  |  |  | X |
| GO:0097305 | response to alcohol | BP | X | X | X | X | X | X | X |
| GO:0009737 | response to abscisic acid | BP |  | X | X | X | X | X | X |
| GO:0071229 | cellular response to acid chemical | BP |  | X | X |  |  |  | X |
| GO:0042631 | cellular response to water deprivation | BP |  | X | X |  |  |  | X |
| GO:0071462 | cellular response to water stimulus | BP |  | X | X |  |  |  | X |
| GO:0019853 | L-ascorbic acid biosynthetic process | BP | X |  | X |  |  |  | X |
| GO:0009963 | positive regulation of flavonoid biosynthetic process | BP |  |  |  |  | X |  | X |
| GO:0019632 | shikimate metabolic process | BP |  |  |  |  |  | X | X |
| GO:0016772 | transferase activity, transferring phosphorus-containing groups | MF | X |  |  | X |  | X | X |
| GO:0016301 | kinase activity | MF | X |  |  | X |  | X | X |
| GO:0016773 | phosphotransferase activity, alcohol group as acceptor | MF | X |  | X | X |  | X | X |
| GO:0004672 | protein kinase activity | MF | X |  | X | X |  | X | X |
| GO:0004674 | protein serine/threonine kinase activity | MF | X |  | X | X |  | X | X |
| GO:0106310 | protein serine kinase activity | MF | X |  | X | X |  | X | X |
| GO:0106311 | protein threonine kinase activity | MF | X |  | X | X |  | X | X |
| GO:0050660 | flavin adenine dinucleotide binding | MF |  |  |  | X |  |  | X |
| GO:0000287 | magnesium ion binding | MF |  |  |  |  | X |  | X |
| GO:0015144 | carbohydrate transmembrane transporter activity | MF |  |  |  |  | X |  | X |
| GO:0090599 | alpha-glucosidase activity | MF |  | X | X |  |  |  | X |
| GO:0003855 | 3-dehydroquinate dehydratase activity | MF |  |  | X |  |  | X | X |
| GO:0004764 | shikimate 3-dehydrogenase (NADP+) activity | MF |  |  | X |  |  | X | X |
| GO:0008143 | poly(A) binding | MF |  |  |  |  |  | X | X |
| GO:0070717 | poly-purine tract binding | MF |  |  |  |  |  | X | X |
| GO:0030136 | clathrin-coated vesicle | CC |  |  | X |  | X | X | X |
| GO:0031312 | extrinsic component of organelle membrane | CC |  |  |  |  | X | X | X |
| GO:0031314 | extrinsic component of mitochondrial inner membrane | CC |  |  |  |  | X | X | X |
| GO:0012510 | trans-Golgi network transport vesicle membrane | CC |  |  |  |  | X |  | X |

## DISCUSSION

In this study, we characterized the DNA methylation response to a panel of six different environmental treatments in the clonal tree *Populus nigra cv.* ‘Italica’. To the best of our knowledge, this is the first WGBS study that evaluates the effects of many different environmental factors on a tree species, and with a high number of replicates (n=8), thereby enabling a robust and comprehensive comparative analysis of the DNA methylation stress response.

### Global signatures of the poplar methylome response to individual stress treatments

#### Abiotic stresses

The global patterns of DNA hypermethylation after exposure to abiotic stresses substantiates previous studies on Populus that have reported global DNA methylation increases after 5-weeks of drought stress in *P. trichocarpa* (Liang et al., 2014) and after 7 days of salt stress in *P. euphratica* (Su et al., 2018). In *P. simonii*, methylation gradually increased during the first 24 h of either cold, heat, salinity or osmotic stress, and certain enzymes involved in (de)methylation pathways were up/downregulated in a stress-specific manner (Song et al., 2016). However, the low-resolution methods (HPLC and MSAP) used to quantify DNA methylation did not allow the authors of that study to further investigate methylation at context-specific level. Here, using WGBS data, we were able to determine that genome-wide stress-induced hypermethylation can arise in a sequence context specific manner as a response to specific stresses. Together with the mentioned studies, our findings suggest that the context-specific hypermethylation patterns induced by abiotic stresses are the product of *de novo* methylation via the RdDM pathway combined with stress-specific up/downregulation of demethylation pathways.

Abscisic acid (ABA) is known to initiate stress signalling leading to physiological acclimation upon stress (Jia et al., 2016, 2017; Popko et al., 2010). However, only few studies have hinted its potential role in mediating global hypermethylation responses (Lafon-Placette et al., 2018; Song et al., 2016; Su et al., 2018). For instance, ABA treatments in Arabidopsis induced hypermethylation at ABA-responsive genes (Gohlke et al., 2013). Moreover, ABA-mediated upregulation of specific microRNAs can downregulate targeted demethylases (Sunkar & Zhu, 2004), which in turn may result in hypermethylation. Because increase of methylation may be associated with gene silencing (Fojtova et al., 2003; Paszkowski & Whitham, 2001), stress-induced global hypermethylation may induce progressive gene silencing leading to arrested growth under adverse conditions. Conversely, In *P. tremula* and tree peony, bud growth reactivation is preceded by a progressive reduction of genomic DNA methylation (Conde et al., 2017; Y. Zhang et al., 2020). Therefore, it is plausible that cold-induced hypermethylation occurs as a first response during winter, arresting growth, followed by a gradual demethylation that leads to growth reactivation in spring. Since changes in CHH methylation are less stable than those in CpG/CHG contexts (Secco et al., 2015; Wibowo et al., 2016), context-specific hypermethylation suggests different stabilities and thus durations of the response. This might reflect differences in duration of the environmental stresses in nature, specifically longer cold periods (winter) versus brief episodes of drought during the growing season.

#### Biotic stresses

The effect of biotic stresses on DNA methylation has been examined in *Arabidopsis* and other species (Dowen et al., 2012; Ramos-Cruz et al., 2021; H. Zhang et al., 2018), however little information is available about woody plants. It is known that SA treatment, rust infection, and caterpillar attack increase the levels of SA, JA and ABA in the affected poplar leaves (Clavijo Mccormick et al., 2014; Eberl, Hammerbacher, et al., 2018; Eberl, Perreca, et al., 2018; Li et al., 2018; Ullah et al., 2019). Moreover, SA can be transported from infected to uninfected sites to induce systemic acquired resistance (SAR) (Li et al., 2018). Therefore, we will discuss the biotic-induced methylation patterns in the context of SAR as we sampled non-directly affected leaves that grew during the stress periods.

Consistent with our results, treatment with exogenous SA has been reported to induce DNA hypomethylation in other species, which in turn activates defense-response genes (Dowen et al., 2012; Ngom et al., 2017). Specifically, in *Vitis amurensis*, exogenous SA upregulated specific demethylases (Kiselev et al., 2013), which induced hypomethylation and consequently enhanced production of secondary metabolites (Kiselev et al., 2015). The loss of DNA methylation can either prime (upon removal of CHH methylation) or constitutively derepress (upon removal of CpG/CHG/CHH methylation) the SA-dependent defense response (Deleris et al., 2016; López Sánchez et al., 2016). Since we found similar CpG/CHG hypomethylation patterns upon both herbivory and SA treatment, we speculate that such response mainly de-repress the SA-dependent defense response, while the TE-associated CHH hypomethylation response after rust infection could have a priming effect.

Diminished rust infection has been observed in drought-affected poplars, explained to some extent by increased stomatal closure mediated by ABA (Ullah et al., 2019), but disregarding the interplay between methylation responses. Here, we showed that rust infection also induced CpG/CHG hypermethylation profiles very similar to those observed under drought and heat, but not cold. These similarities suggest a possible overlap between the responses to drought and rust infection, as it has been suggested by results on the poplar apoplast proteome (Pechanova et al., 2010).

### Hotspots of environmental-induced methylation variation

Experiments for studying stress effects on DNA methylation usually analyse DMRs induced by single stresses. Thus, intersection of several DMR sets detected from different stresses allows the capture of more generic responses. Though the latter approach can pinpoint multi-stress DMRs (Song et al., 2016; Xue et al., 2013), it likely underestimates commonalities in the methylation response to different stresses because stringent significance thresholds in DMR detection can leave most of the responding loci undetected. Here, we found that CpG/CHG-DMRs often showed a similar response irrespective of treatment, suggesting that much of the stress response in poplar is generic, rather than stress-specific. Based on these observations, we suspect that many of the reported stress-specific DMRs in other species likely have also a multi-stress nature, which would imply a more careful interpretation of DMR results in the future.

Our results resemble the observations of epimutational hotspots in nearly isogenic *Arabidopsis* lines under greenhouse and natural environmental conditions (Becker et al., 2011; Hagmann et al., 2015; Schmitz et al., 2011). Such epimutation hotspots are characterized by steady-state intermediate methylation levels (Hazarika et al., 2022), which was also observed in the control methylation levels of CG/CHG DMRs. However, where the intermediate methylation level of *A. thaliana* hotspots is due to sparse cytosine methylation (only a subset of CpGs is methylated), in our case it is the result of individual cytosines being partially methylated. Since such CpG/CHG-DMRs are often located on TE flanking regions, we hypothesize that TE-mediated stress-induced (de)methylation is the source of methylation variation on the TE edges, here identified as multi-stress DMRs.

### Stress-induced methylation variation as a source of epialleles under natural conditions

In this clonal system, the accumulation of methylation variation can be attributed mostly to spontaneous and environmentally induced variation. In other species, transient stress-responsive epigenetically labile regions have been identified to also overlap with naturally occurring DMRs, suggesting a non-random stress-triggered epigenetic reprogramming (Miryeganeh et al., 2022; Wibowo et al., 2016). Here, we identified many CpG/CHG-DMRs among ortets, which are thought to be mitotically stable and hence clonally transmissible. More interestingly, a large proportion of stress-induced DMRs overlapped with ortet-DMRs. This result indicates that at least part of the natural methylation variation of the clonal system at a European scale is induced by changing environments. Consequently, environment-induced methylation variants in CpG/CHG contexts could be fixed and appear as natural epialleles detectable across the tree lifespan and maybe next clonal generations.

In contrast, induced CHH-DMRs showed only a minor overlap with ortet-DMRs, even though such DMRs largely arose in response to drought and heat. This observation supports the idea that CHH methylation variation quickly disappears after the stress is gone, preventing induced CHH-DMRs to persist as natural epialleles. This capability of CpG/CHG methylation to track long-term environmental variation seems to be supported by recent observations in other trees (Heer et al., 2018; Miryeganeh et al., 2022).

### Functionality of the poplar methylome response to drought

As poplar is a fast-growing riparian tree whose high productivity requires high water availability (Monclus et al., 2006; Vanden Broeck, 2003), methylation responses to drought are potentially relevant for the ecology of this species. Recent reports in the species have found significant genotypic variation involved in drought tolerance (Viger et al., 2016) as well as for drought escape (Yıldırım & Kaya, 2017). Therefore, efficient finetuning of the drought escape and tolerance responses is likely a strong selection pressure in this clonal cultivar, which may have promoted the evolution of DNA-methylation-based regulatory mechanisms.

Even though the study of the functionality of DNA methylation would require at the very least quantification of gene expression, some patterns that we observed in the methylome response to drought suggest a functional consequence. Here, we reported TE-associated CHH hypermethylation mostly in gene flanking regions, which has been also described in *P. trichocarpa* (Liang et al., 2014). However, our analysis also revealed hypermethylation enriched on specific TE superfamilies: SINE and MITE/DTH.

TE activity can be triggered by biotic and abiotic stress conditions (Lanciano & Mirouze, 2018; Seibt et al., 2016), which in turn may lead to a rapid and extensive TE amplification followed by inactivity and drift (Jiang et al., 2004). Such is the case of SINEs and MITEs irrespective of their inherent differences: retrotransposons vs. DNA transposons, respectively. Both superfamilies are relatively short elements frequently inserted close to and within genes (Kögler et al., 2020; Seibt et al., 2016), likely due to their tendency to integrate in hypomethylated DNA regions (Arnaud et al., 2000). Also, both are preferential targets for de novo methylation, which can then spread into flanking sequences and may affect the expression of nearby genes (Arnaud et al., 2000; Chen et al., 2014). TE proximity to genes may suggest that stress-induced TE hypermethylation could be a by-product of highly expressed nearby genes, as previously suggested by (Secco et al., 2015). However, we observed that CHH hypermethylation occurred irrespective of their distance to genes, indicating that such response may not be a consequence of nearby gene expression.

Hypermethylation of entire TE superfamilies in response to stress has not been previously reported in other species. We found that SINE and MITE/DTH elements were already highly methylated (and presumably silenced) under control conditions. Therefore, as drought seems to reinforce such hypermethylation, we speculate that the selective silencing of these elements can have regulation consequences of nearby drought-responsive genes as hinted by GO enrichment results.

Genes associated with drought CHH-DMRs seemed enriched in general responses to drought, such as ABA signalling, protein kinase activity, and response to water deprivation. Remarkably, genes associated with SINE and MITE/DTH elements were also enriched in similar functional responses. Evidence of MITE-derived small RNAs regulating abscisic acid signalling and abiotic stress responses in rice (Yan et al., 2011) suggests that these elements may have been selected and retained close to specific genes that play a role in the rapid response to drought. Therefore, we speculate that the triggered hypermethylation response is regulating, via specific TE superfamilies, genes and pathways involved in functional responses to drought. To test this functional hypothesis, it will be important in follow-up studies to monitor expression and methylation of SINEs, MITE/DTHs, and the nearby genes prior to and during drought stress.

In conclusion, our evaluation of the poplar methylome plasticity upon abiotic and biotic treatments allowed the discovery of multi-stress hotspots that are partially shaping the natural methylation variation. Moreover, we were able to identify specific TE superfamilies whose response to drought may have been selected to cope with extreme conditions. Our study furthermore highlights the importance of analysing the effect of multiple factors in the same experiment to avoid overstatements of individual effects.

## DATA AVAILABILITY

The bisulfite sequencing raw data is deposited in the ENA Sequence Read Archive Repository (www.ebi.ac.uk/ena/) under study accession number: PRJEB51831.

Methylation files for the three contexts and the list of annotated differentially methylated regions are deposited in zenodo (https://zenodo.org/) under DOI: 10.5281/zenodo.7193978

Gene models and TE predictions for the poplar clonal cultivar used in this study are deposited in zenodo under DOI: (to be uploaded)

## ACKNOWLEDGEMENTS

We thank Slavica Milanovic-Ivanovic and Gregor Disveld for their technical help in the molecular lab and greenhouse settings, respectively. We thank Morgane van Antro and Haymanti Saha for the fruitful discussions and sampling assistance. We thank Bhumika Dubay for her work on the reference genome, gene model and TE predictions. We are grateful for all the input and discussions with all the members of the Epidiverse Consortium. We also express our gratitude to Sybille Unsicker for providing the poplar rust spores and gypsy moth caterpillars.

This work was supported by the European Training Network “EpiDiverse” and received funding from the EU Horizon 2020 program under Marie SkłodowskaCurie grant agreement No 764965.

## AUTHOR CONTRIBUTIONS

**Peña-Ponton C.:** Conceptualization, methodology, software, formal analysis, investigation, data curation, writing - original draft, visualization

**Díez-Rodríguez B.:** Resources, writing - review & editing

**Perez-Bello P.:** Resources, writing - review & editing

**Becker C.:** Investigation, writing - review & editing

**McIntyre LM.:** Formal analysis, writing - review & editing

**Van der Putten. W.:** Writing - review & editing, supervision

**De Paoli E.:** Conceptualization, writing - review & editing

**Heer K.:** Conceptualization, resources, writing - review & editing, supervision, project administration, funding acquisition

**Opgenoorth L.:** Conceptualization, resources, writing - review & editing, supervision, project administration, funding acquisition

**Verhoeven KJF.:** Conceptualization, formal analysis, writing - original draft, writing - review & editing, supervision, project administration, funding acquisition

