## Supporting information for "High-resolution methylome analysis in the clonal *Populus nigra* cv. ‘Italica’ reveals environmentally sensitive hotspots and drought-responsive TE superfamilies"

### *New Phytologist* Supporting Information

Article acceptance date: Click here to enter a date.

The following Supporting Information is available for this article:

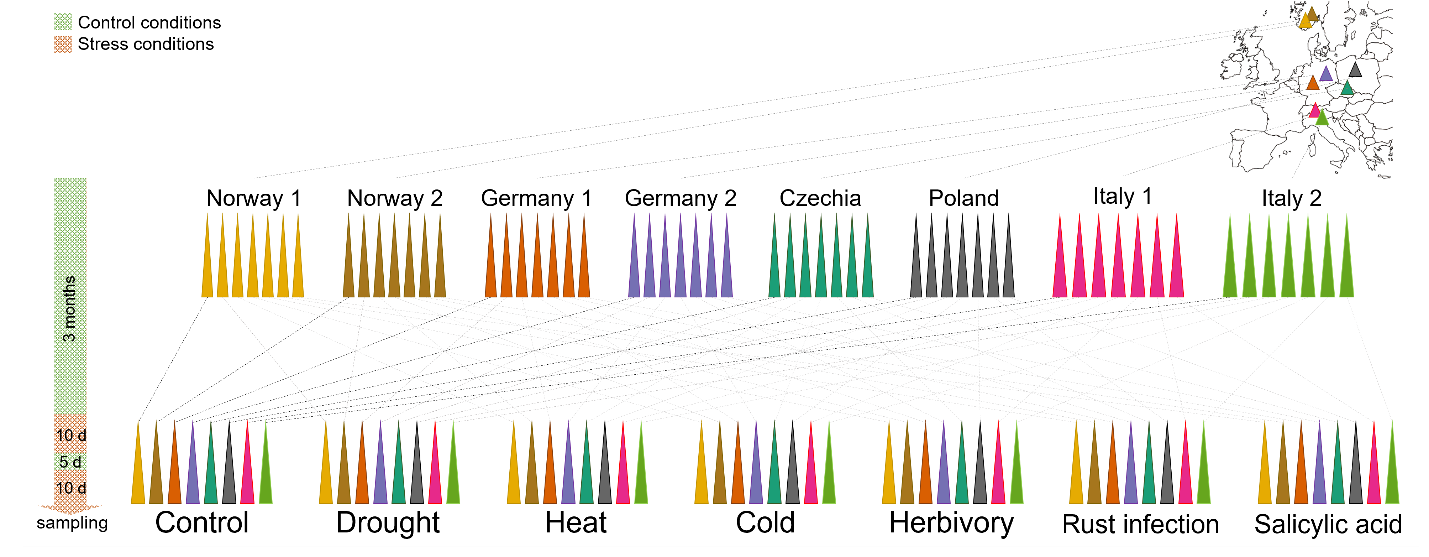

**Fig. S1** Stress experimental design. Eight adult poplar trees in the field (ortets) were clonally propagated into at least seven ramets per ortet. After 3 months of growth in control conditions*, ramets were exposed to different stress treatments for a total period of 25 days with a recovery period of 5 days in between. At day 26, leaves were sampled for methylation analysis. Different colors represent different ortets. *Temperature: (day/night) 22/18 °C (±2°C), humidity: 60% Rh (±5% Rh), light: (day/night) 16/8 h, VWC: 20.20% (± 3.24 SD)

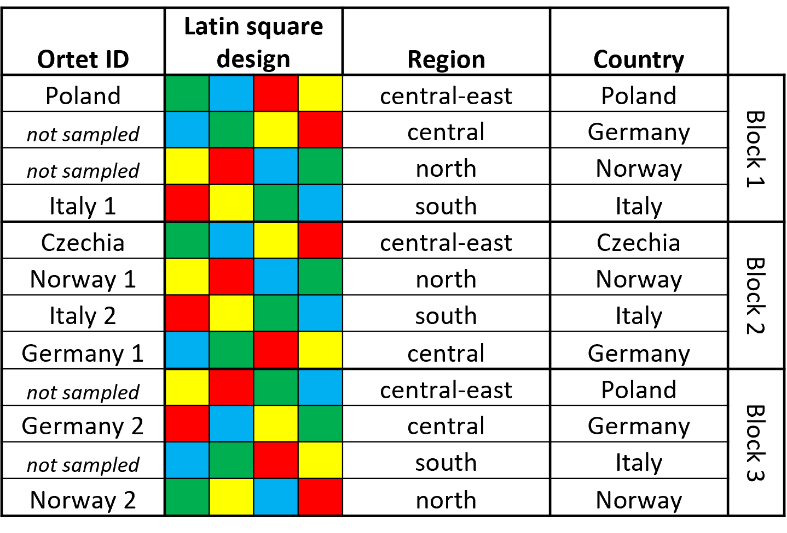

**Fig. S2** Latin square design used for plant allocation on the greenhouse table. Control (green) and drought-treated (yellow) ramets remained on the same positions during the entire experiment, cold-treated (blue) and heat-treated (red) ramets were allocated on the table only during the stress-free period. Three consecutive blocks of Latin squares were arrayed on a long table ensuring that ramets from every geographic region were included on each block. Ramets from 12 ortets were included in the experiment, however after Allegro genotyping (Díez-Rodríguez et al., 2022), only eight were confirmed to belong to the same clonal lineage, which were sampled for the subsequent analyses shown in this publication. Rust infection, herbivory and SA treatments were implemented in separate greenhouses under the same control growth conditions.

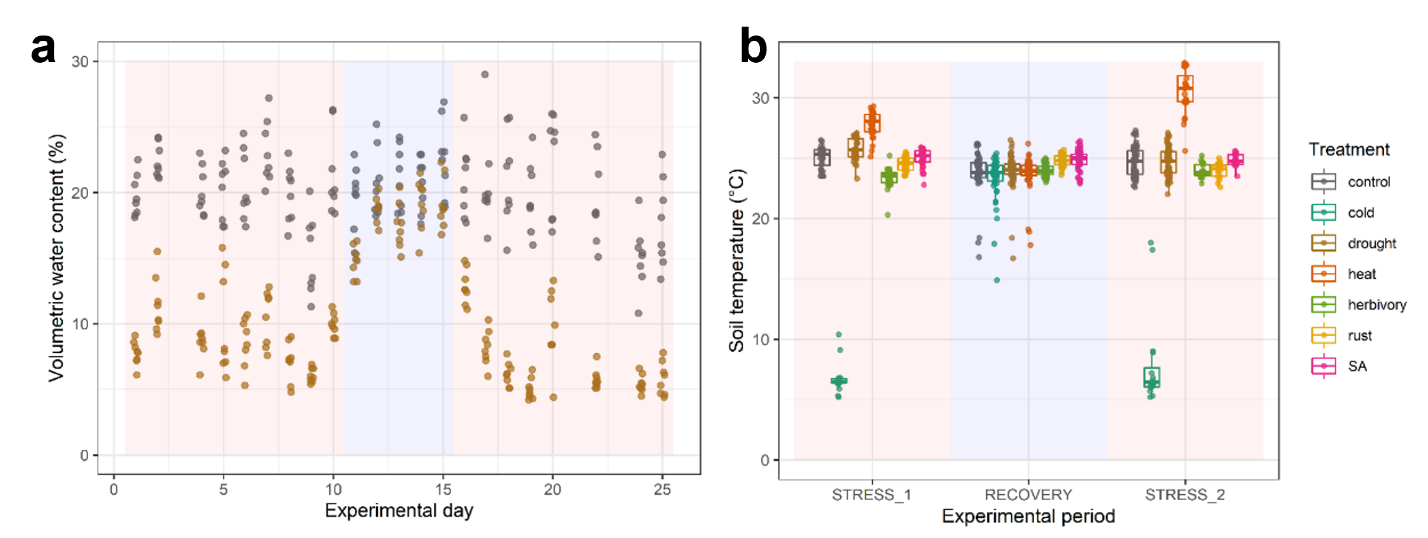

**Fig. S3**  Soil parameters monitored during the experiment. a) Volumetric water content (VWC) is shown for control and drought samples. Two measurements were performed per pot, only the mean VWC is shown. b) Boxplots summarize the soil temperature monitored on each treatment group over the different experimental periods. Stress periods: days 1-10 (STRESS_1), 16-25 (STRESS_2). Recovery periods: days 11-15**.**

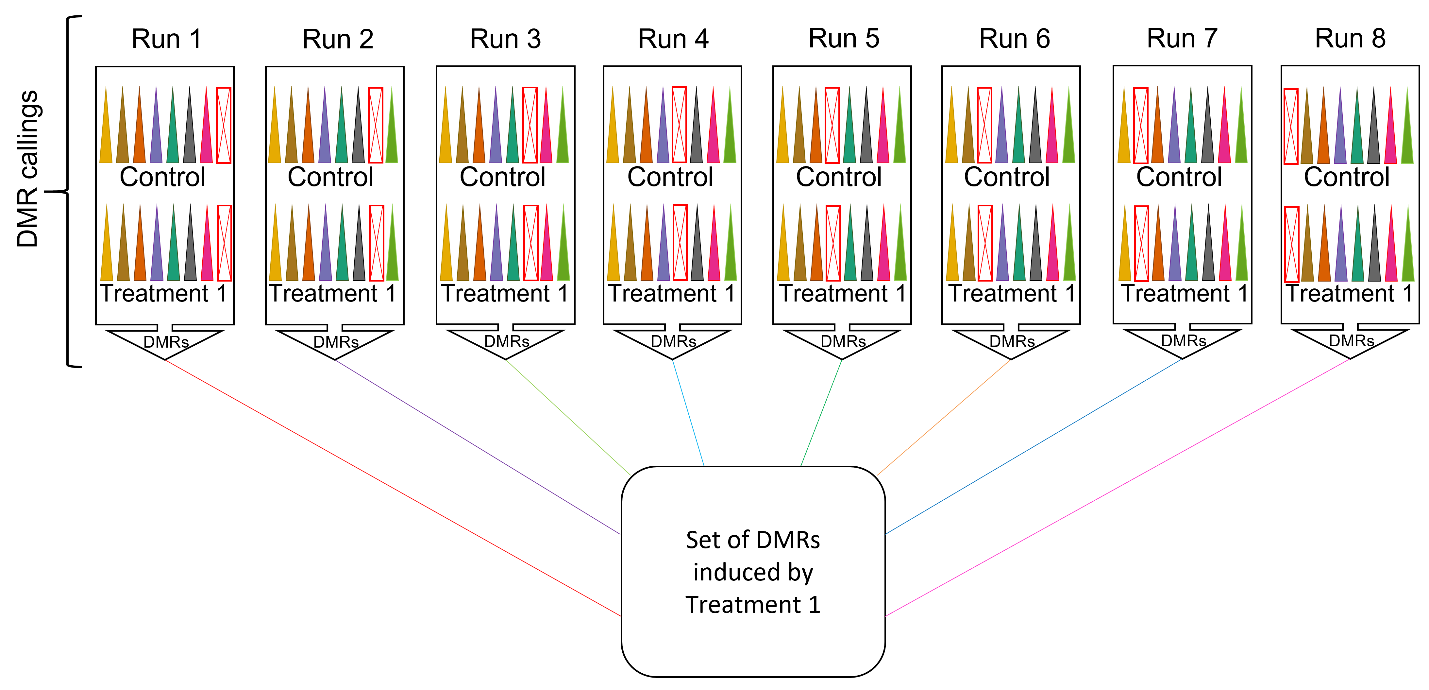

**Fig. S4** Example of DMR calling using jack-knife approach. For each treatment, eight runs were performed leaving one ortet out on each DMR call. All DMR sets were merged for downstream analysis. Different colors represent different ortets. The red crossed rectangles represent the ortet removed from the respective DMR calling.

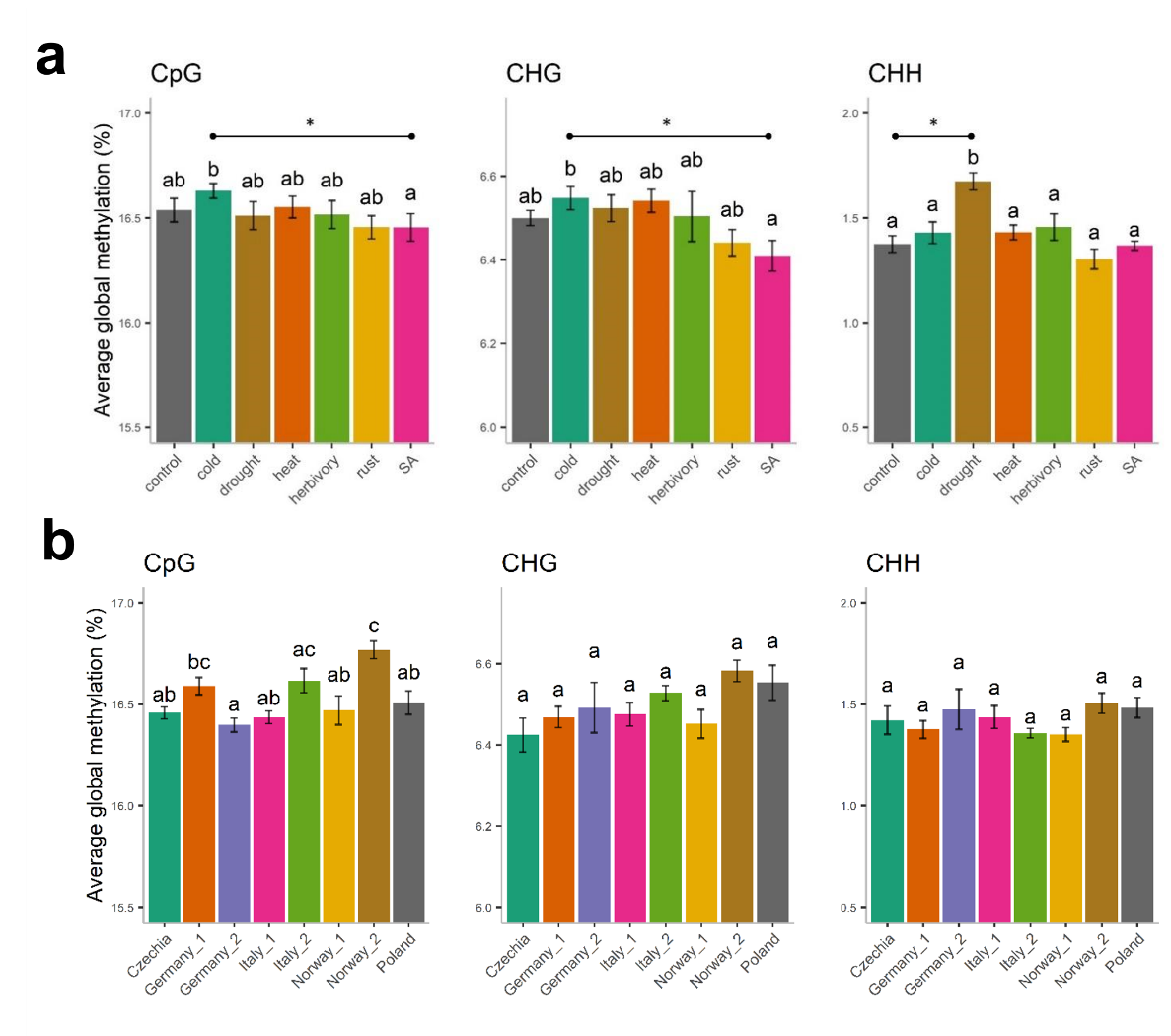

**Fig. S5** Barplots of average global methylation (%) for treatments and ortets a) Comparison between stress treatments, b) comparison between ortets. Sequence contexts were analysed separately. Horizontal bars and letters indicate relevant significant pairwise differences after Tukey post-hoc comparisons.

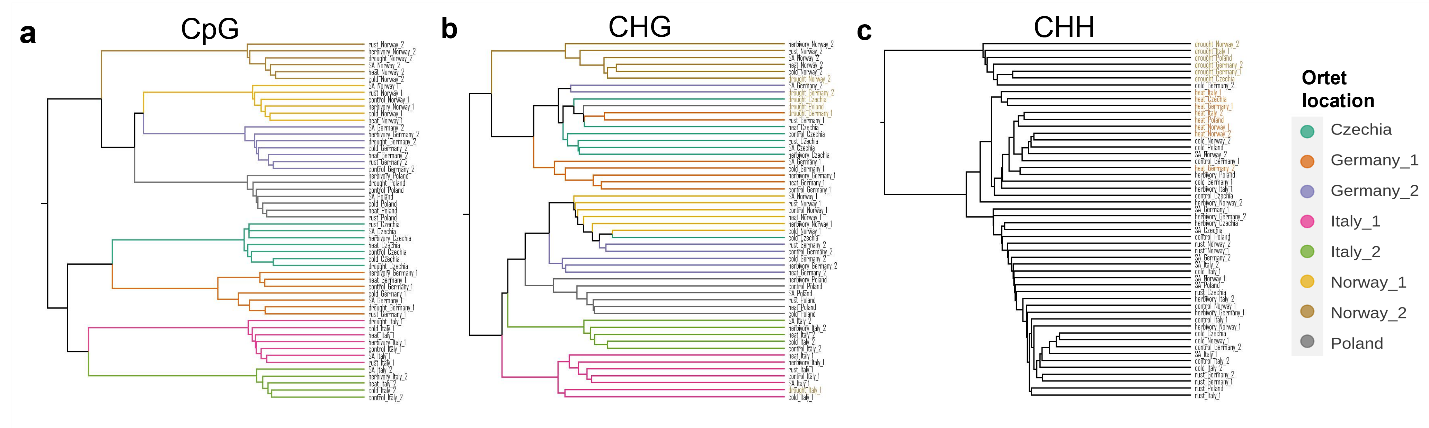

**Fig. S6** Unsupervised hierarchical clustering analysis of CpG (a), CHG (b) and CHH (c) methylation data from stress-treated ramets. In a) and b), dendrogram branches are colored by ortet location, while in c) some representative nodes (individual samples) are colored according to treatment (drought: brown, and heat: orange). For CHH, dendrogram branches were not coloured to highlight colors of the nodes.

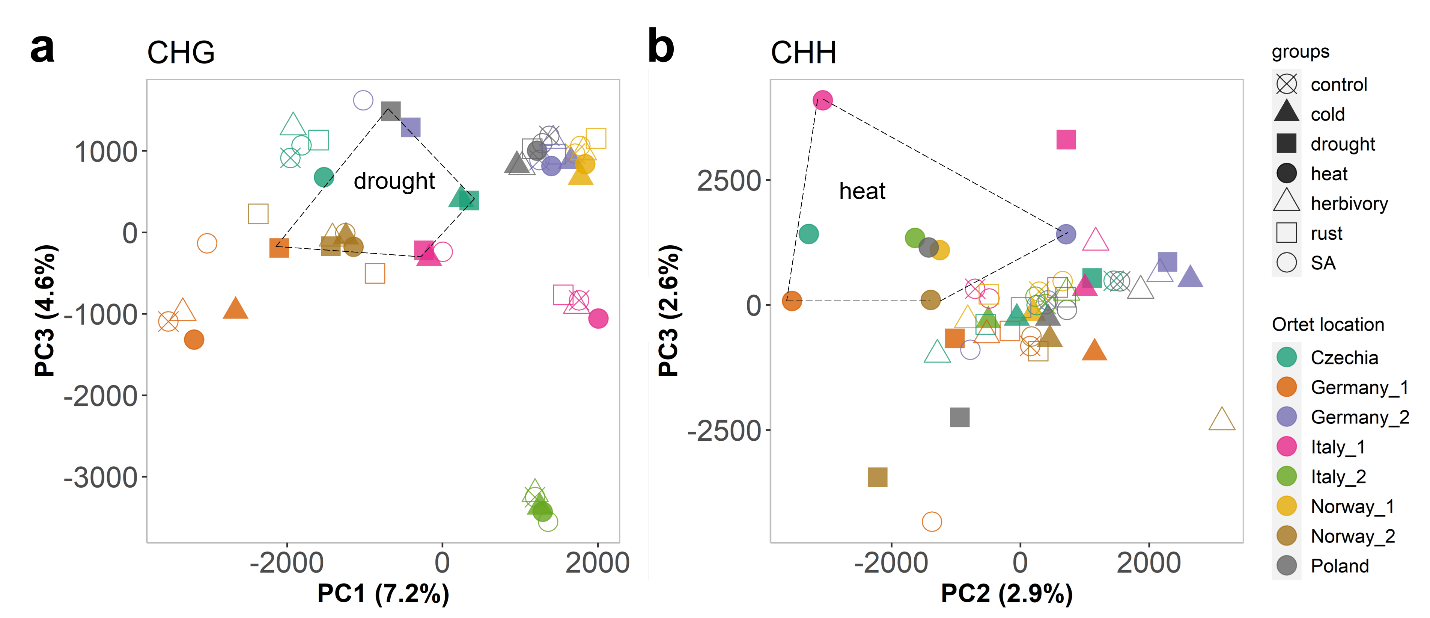

**Fig. S7** Additional informative principal components calculated using methylation data from stress-treated Lombardy poplar ramets. a) PC1 and PC3 are shown for analysis on CHG context. b) PC2 and PC3 are shown for analysis on CHH context. Samples are colored by ortet identity. Different shapes represent each experimental group. Drought and heat clusters are highlighted within dashed lines.

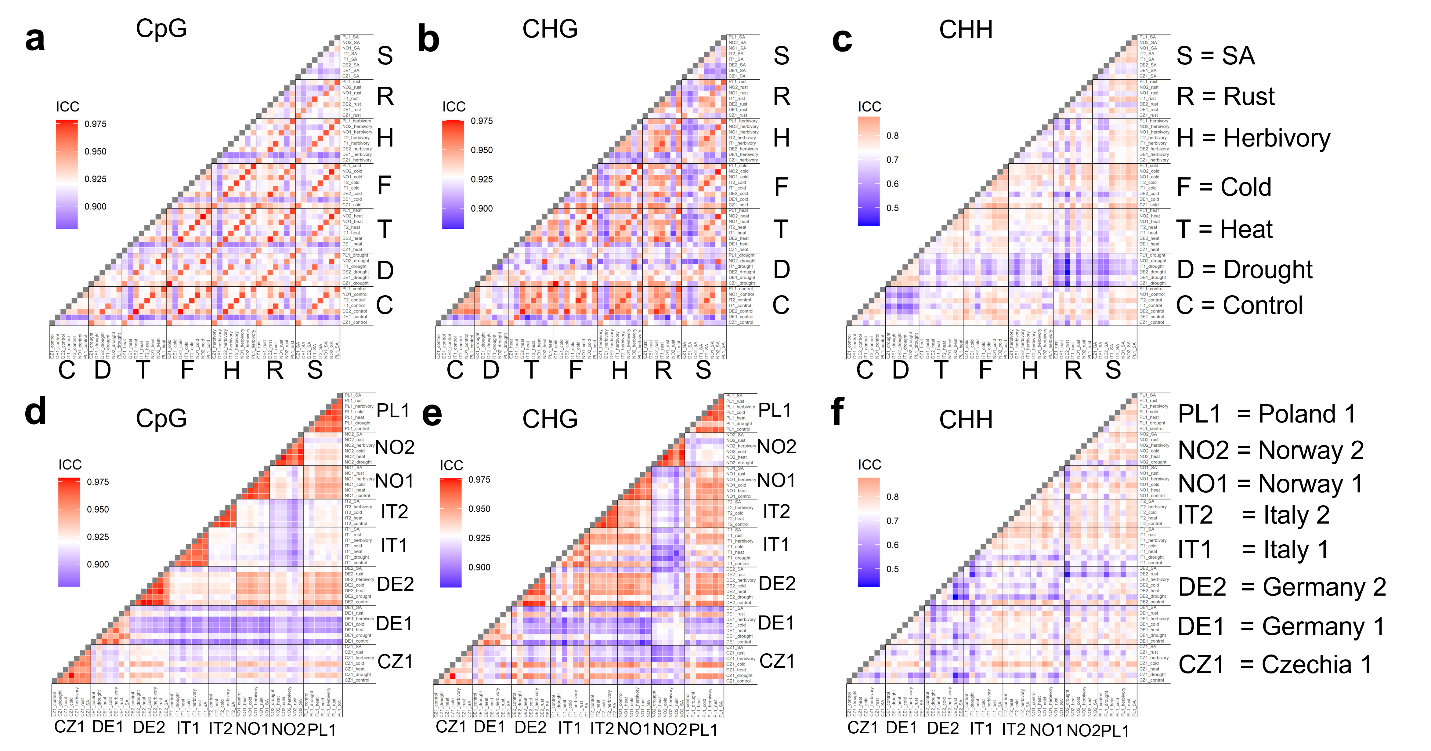

**Fig. S8** Intraclass correlation coefficients (ICC) computed for all ramet pairwise comparisons in the three sequence contexts (CpG, CHG, CHH). On each plot, each coloured tile represent the ICC calculated for the corresponding pairwise comparison. Grids **a)**, **b)**, and **c)** show samples sorted by stress treatment. Grids **c)**, **d)**, and **f)** show samples sorted by ortet location. For each plot, the color gradient was determined by maximum (red), minimum (blue) and median (white) ICC values. The methylation level (%) of 27968, 33779 and 125251 100-bp bins were used for ICC calculations for CpG, CHG and CHH contexts, respectively.

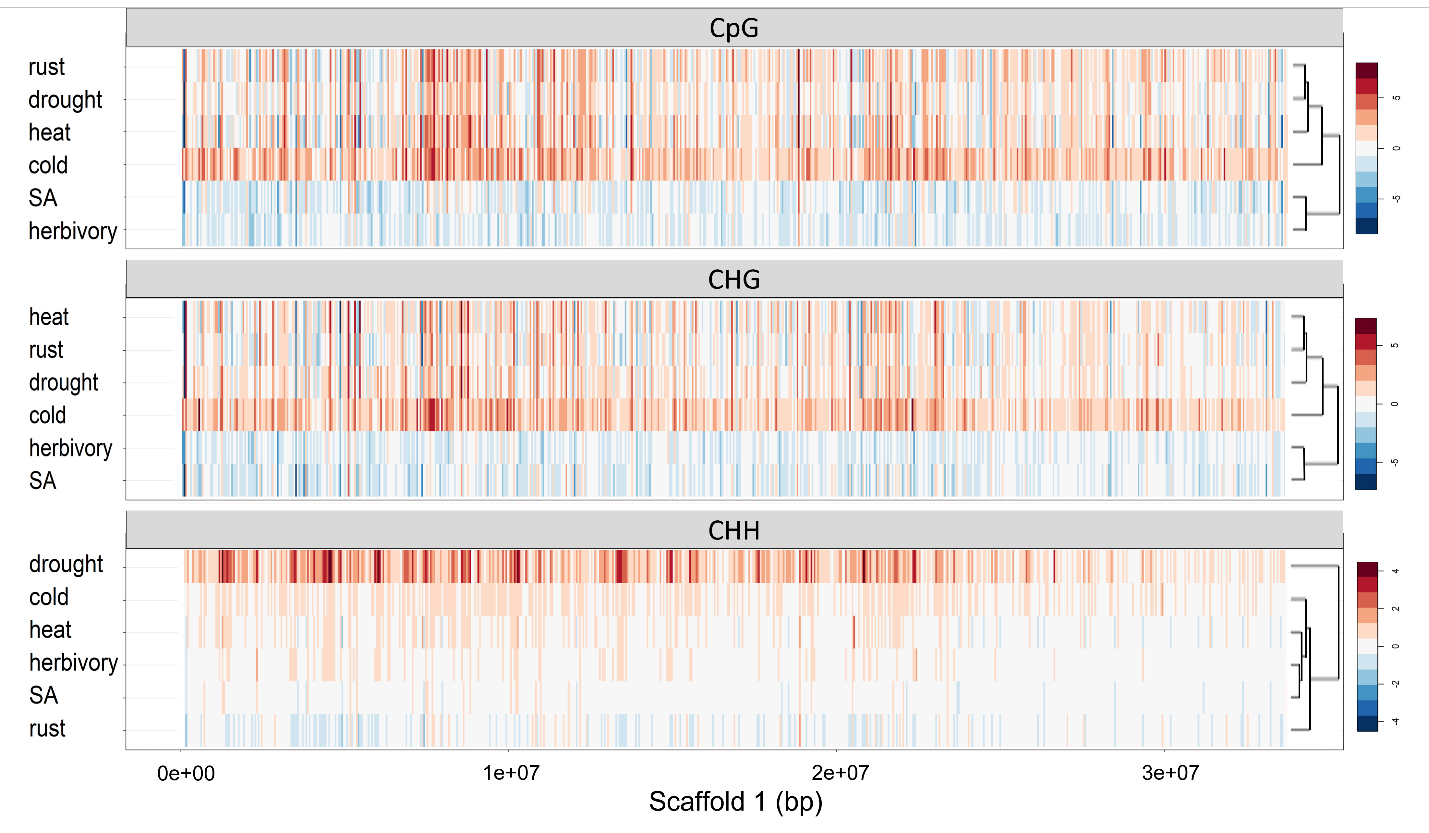

**Fig. S9** Heatmaps of CpG, CHG and CHH methylation variation profiles over the scaffold 1 of *Populus nigra* cv. ‘Italica’. Methylation differences (vs control group) for 50-kb bins were calculated and plotted for each treatment and context. Hierarchical clustering is shown on the right for each context. Methylation differences refer to differences in percentage points

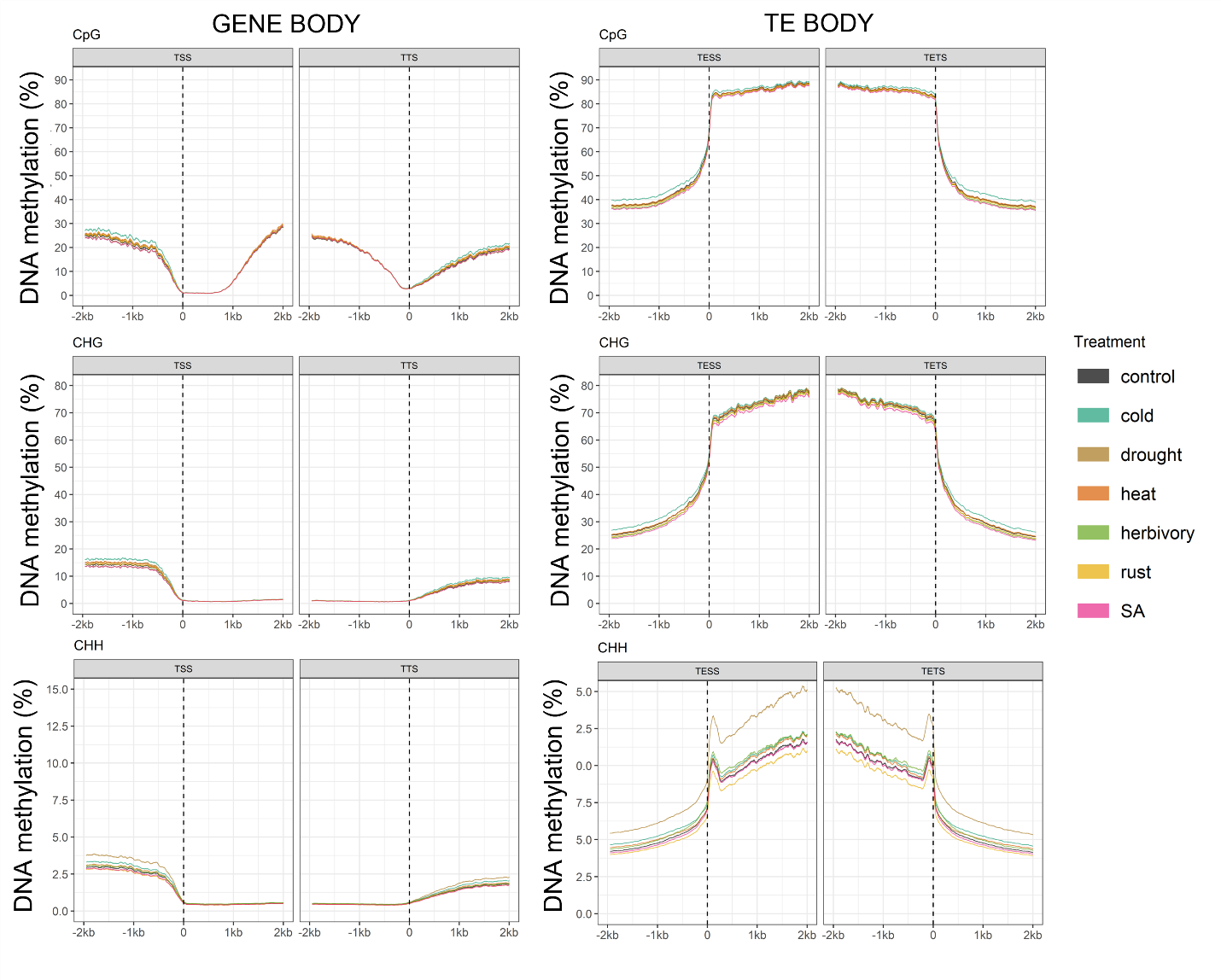

**Fig. S10** Characterization of CpG, CHG and CHH methylation levels within and proximal to gene models and transposable elements. Simple moving averages (SMA) over a period of 50 bp were calculated and plotted for each treatment and context. Left: methylation in gene body and flanking regions. Only protein-coding genes with known 5’UTR and 3’UTR coordinates were analyzed. Right: methylation in TE body and flanking regions. Only >150 bp transposable elements were analyzed. The dashed lines represent the points of alignments of coding-gene transcriptional start site (TSS) or annotated transposable elements start site (TESS) and coding-gene transcription termination site (TTS) or annotated transposable element termination site (TETS). In CHH context, methylation peaks around TESS and TETS correspond to short TEs (~150-200 bp).

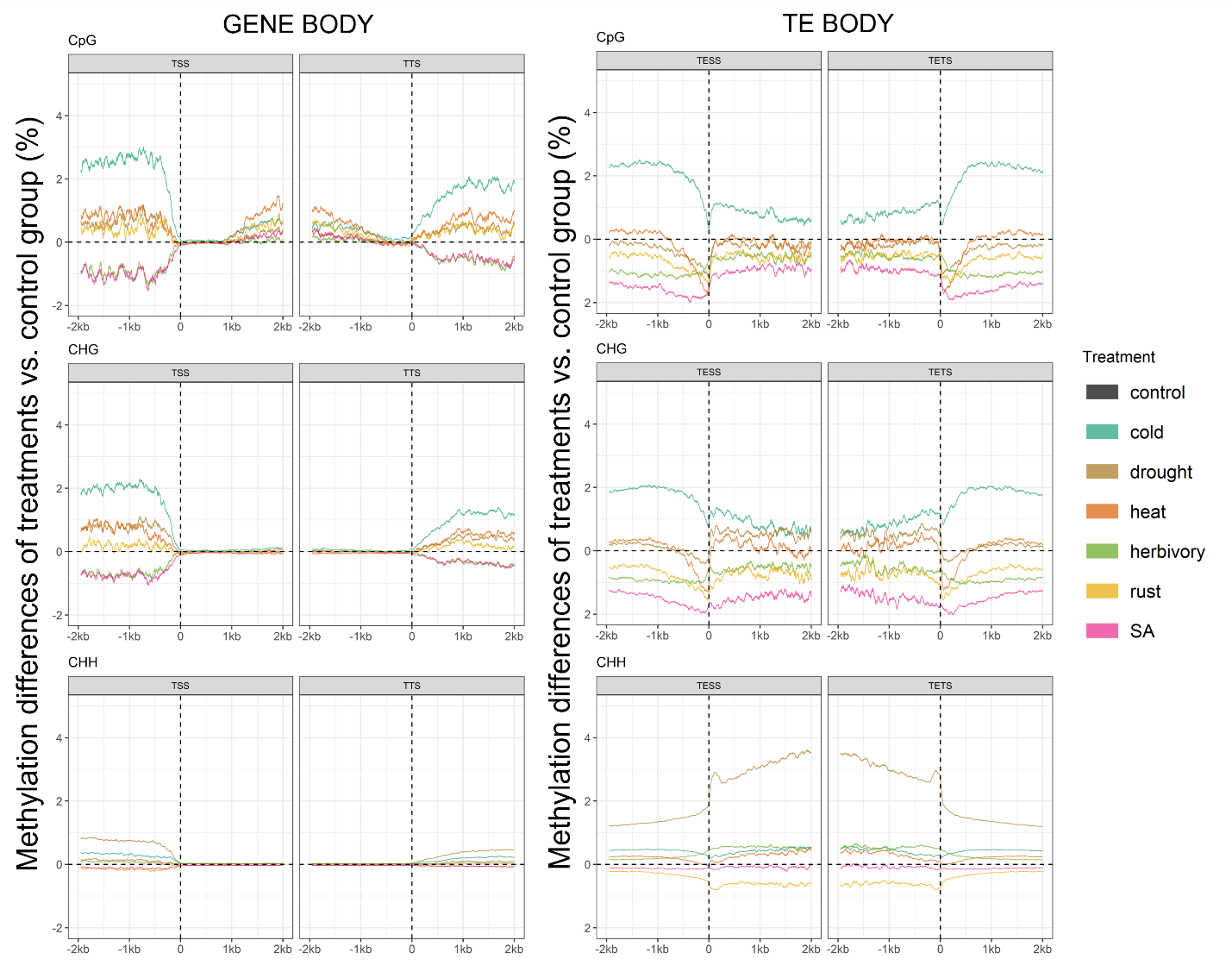

**Fig. S11**  Metaplots of CpG, CHG and CHH methylation level differences (vs control group) within and proximal to gene models and transposable elements. Simple moving averages (SMA) over a period of 50 bp were calculated and plotted for each treatment and context. Left: methylation differences in gene body and flanking regions. Only protein-coding genes with known 5’UTR and 3’UTR coordinates were analyzed. Right: methylation differences in TE body and flanking regions. Only >150 bp transposable elements were analyzed. The vertical dashed lines represent the points of alignments of coding-gene transcriptional start site (TSS) or annotated transposable elements start site (TESS) and coding-gene transcription termination site (TTS) or annotated transposable element termination site (TETS). Methylation differences refer to differences in percentage points

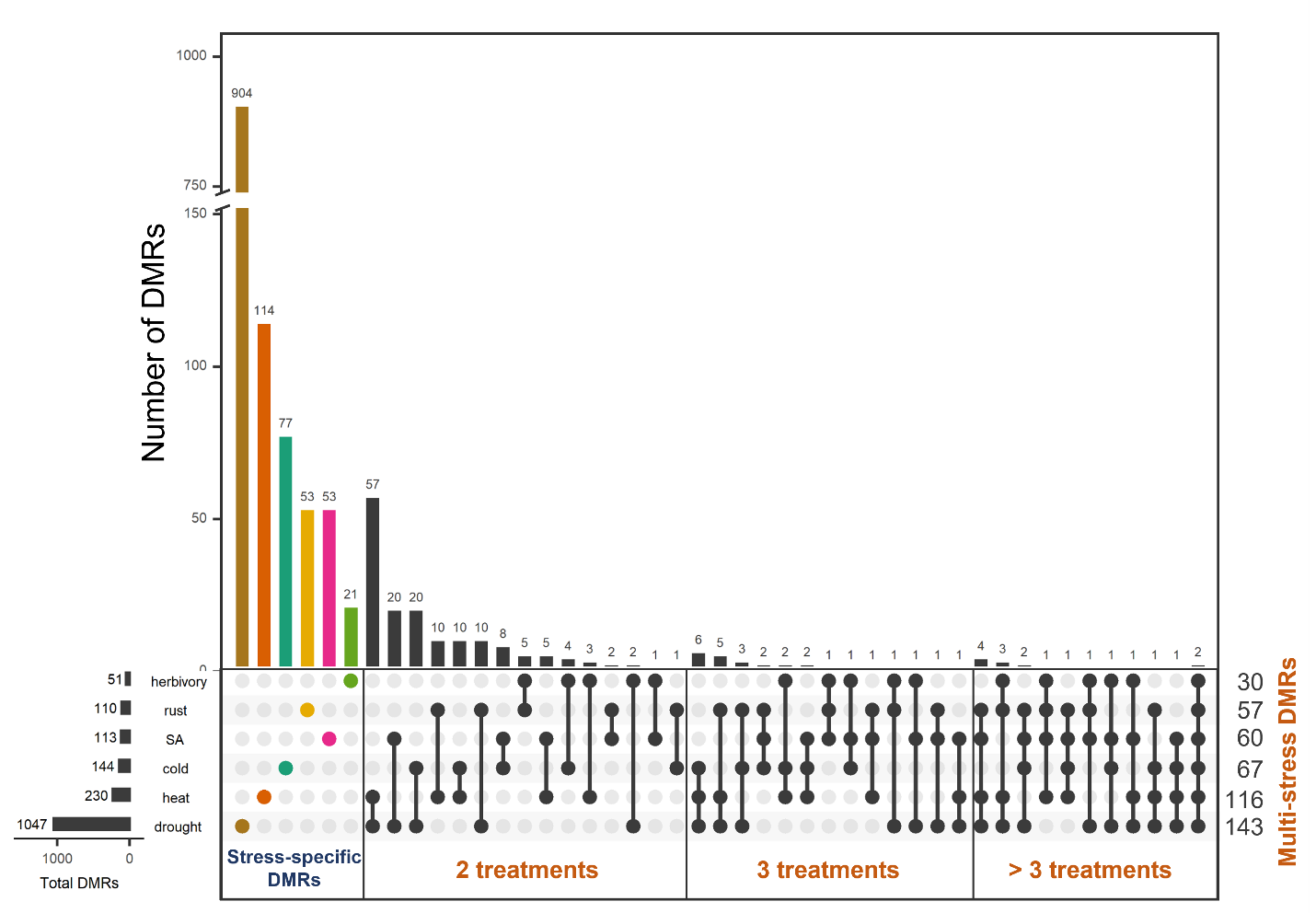

**Fig. S12** Upset plot of DMR set intersections between treatments. Stress-specific DMRs are shown in colored bars and circles. Multi-stress DMRs are shown in black. DMRs in all sequence contexts were grouped together and methylation direction (hyper/hypo) were ignored for the intersections.

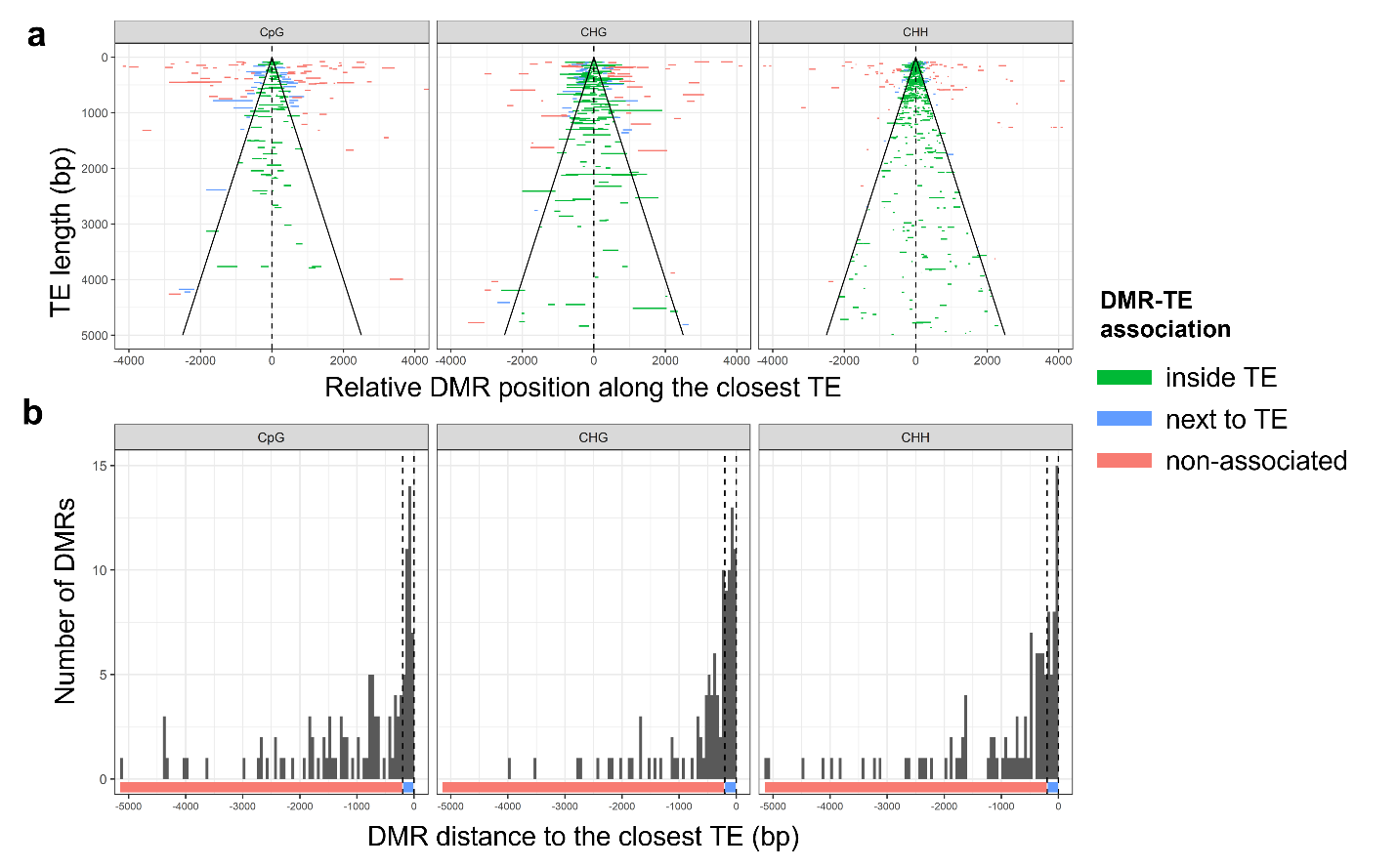

**Fig. S13** DMR association with the closest TE. **a)** DMRs (and DMR length) are plotted according to their relative position to the closest TE. TEs are ordered by length (y axis), the relative DMR positions were calculated based on the corresponding TE midpoint (x axis). Sloping lines depict the TE edges. **b)** Counts of DMRs located close (but not inside) to TEs. DMRs located within the first 200 bp next to a TE were considered as TE-associated. Vertical dashed lines depict the first 200-bp flanking regions of a TE.

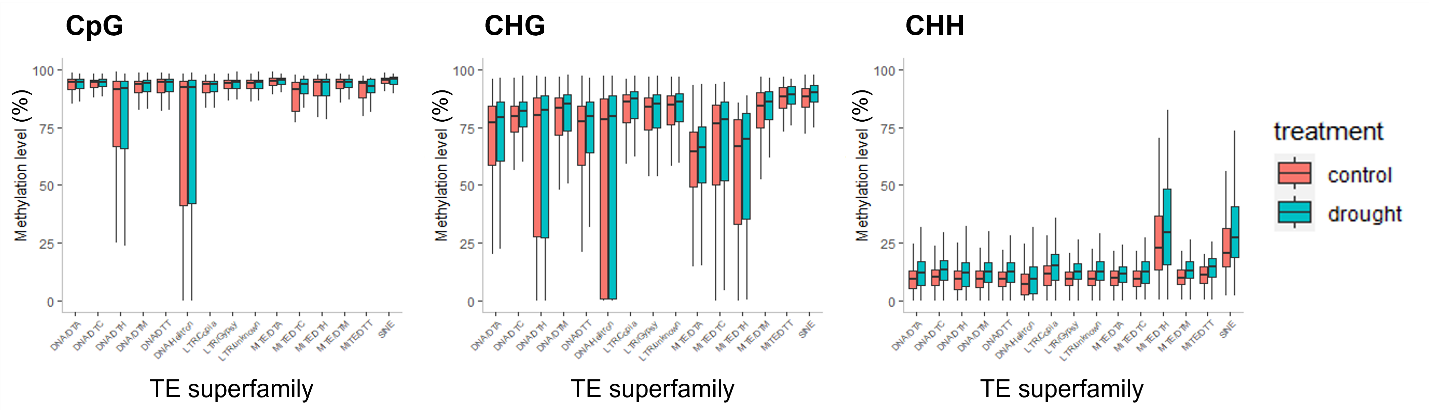

**Fig. S14**  TE methylation levels in all sequence contexts for all TE superfamilies under control and drought conditions.

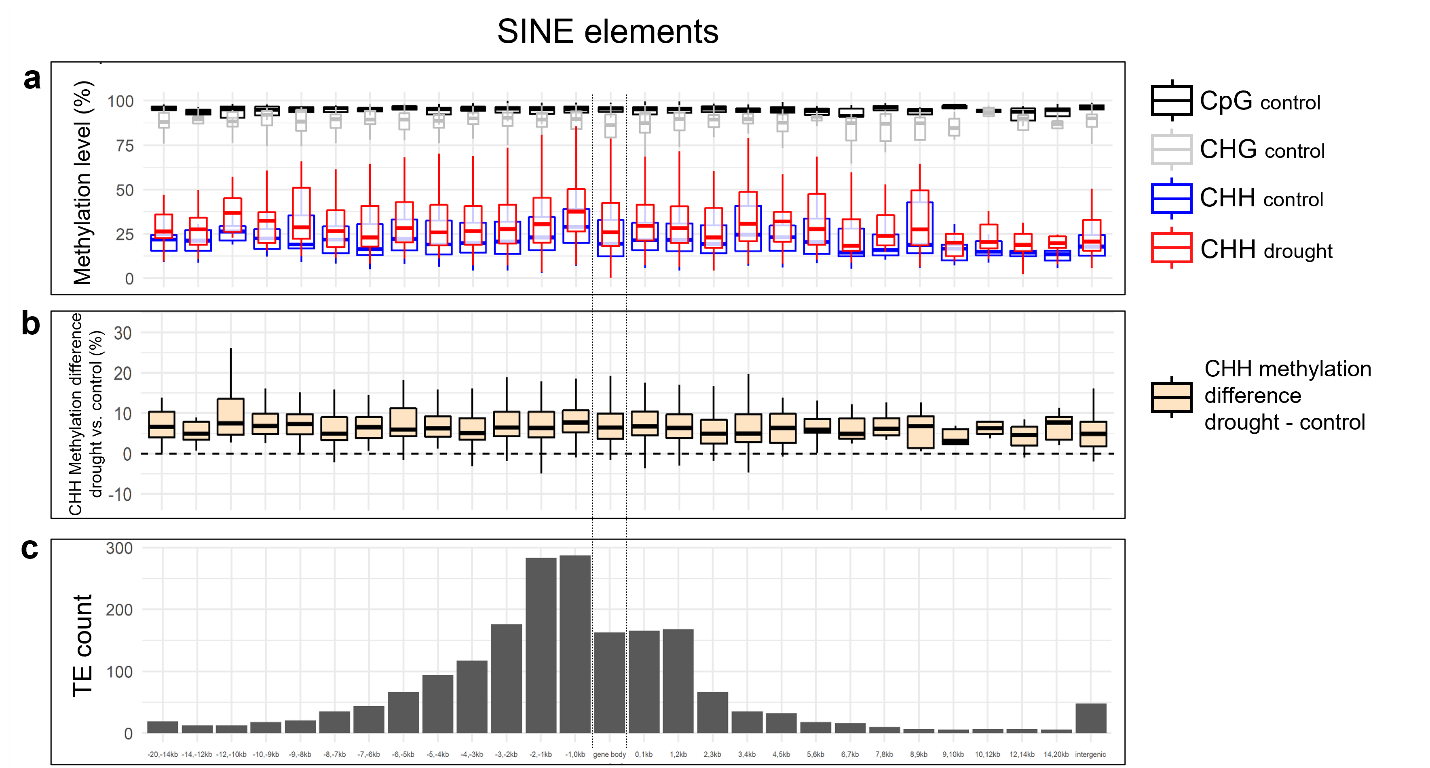

**Fig. S15** Methylation levels for all SINE elements in 1-kb bins along their distribution over genic regions. a) Boxplots of the methylation levels of SINE elements for control CG, CHG and CHH methylation, and drought CHH methylation. b) Boxplots of the difference in CHH methylation levels between SINE elements in drought vs. control group. c) Frequency of SINE elements in 1-kb bins along the genic region. All SINEs located inside gene bodies were included in a single bin (doted vertical lines). Elements located more than 10 kb away from the nearest gene were analyzed in 10-12kb, 12-14kb and 14-20kb bins. Intergenic SINEs (far right) correspond to elements located more than 20 kb away from the nearest gene.

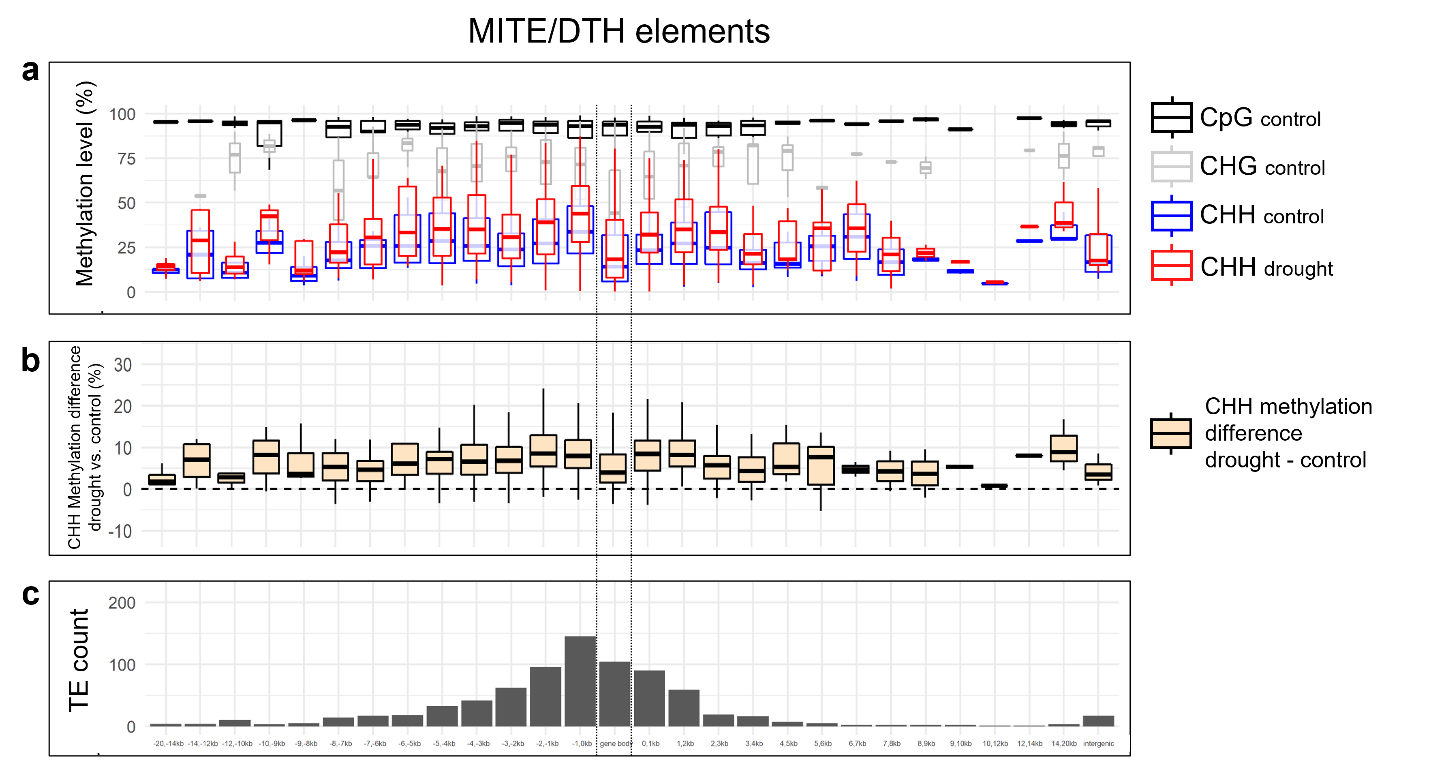

**Fig. S16** Methylation levels for all MITE/DTH elements in 1-kb bins along their distribution over genic regions. a) Boxplots of the methylation levels of MITE/DTH elements for control CG, CHG and CHH methylation, and drought CHH methylation. b) Boxplots of the difference in CHH methylation between MITE/DTH elements in drought vs. control group. c) Frequency of MITE/DTH elements in 1-kb bins along the genic region. All MITE/DTH elements located inside gene bodies were included in a single bin (doted vertical lines). Elements located more than 10 kb away from the nearest gene were analyzed in 10-12kb, 12-14kb and 14-20kb bins. Intergenic MITE/DTHs (far right) correspond to elements located more than 20 kb away from the nearest gene.

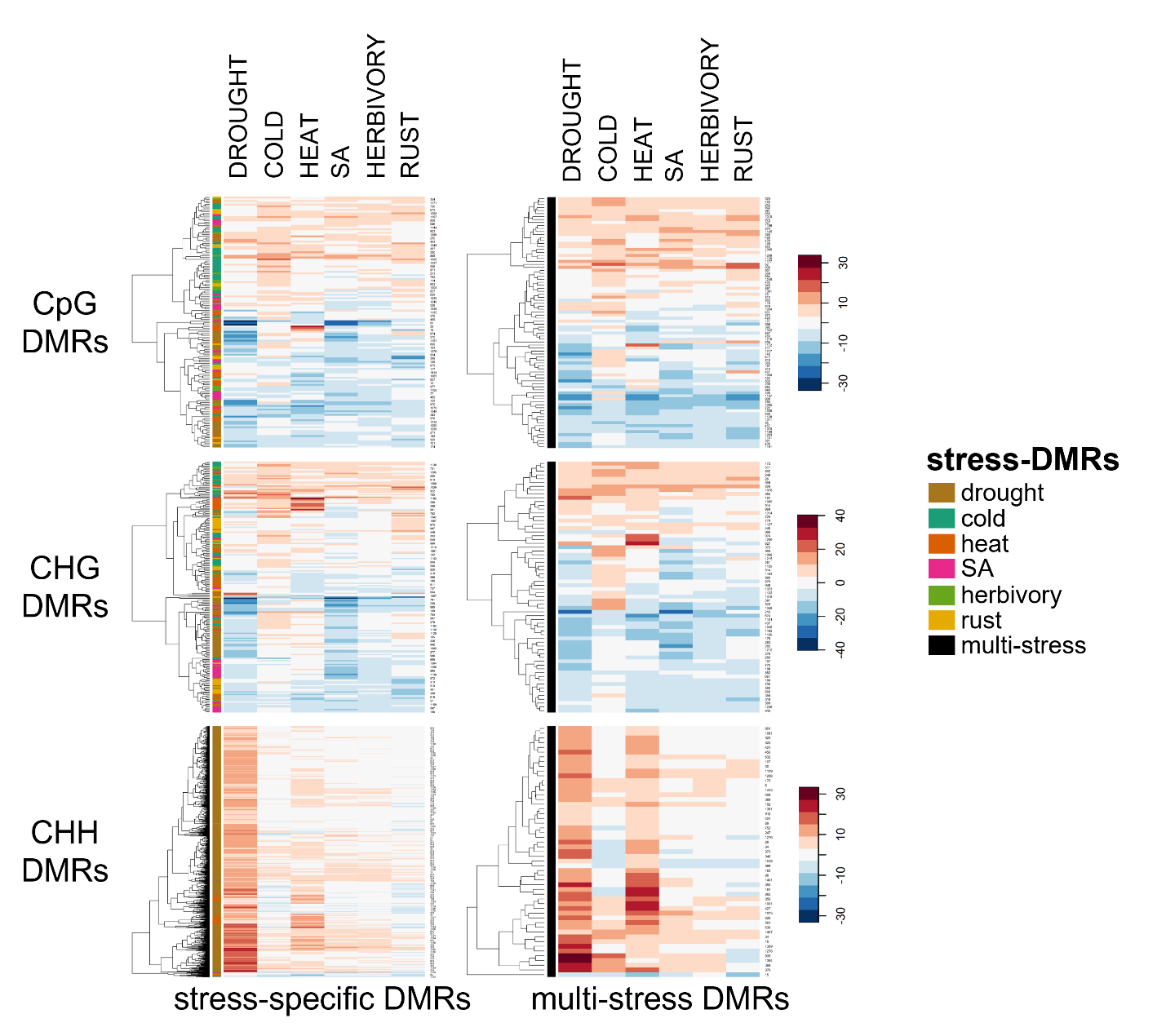

**Fig. S17** Heatmap and hierarchical clustering of the average difference methylation levels (compared to control) of the 1728 identified stress-DMRs in the corresponding sequence context. Left panel: analysis of DMRs identified in only one stress. Right panel: analysis od DMRs identified in more than one stress.

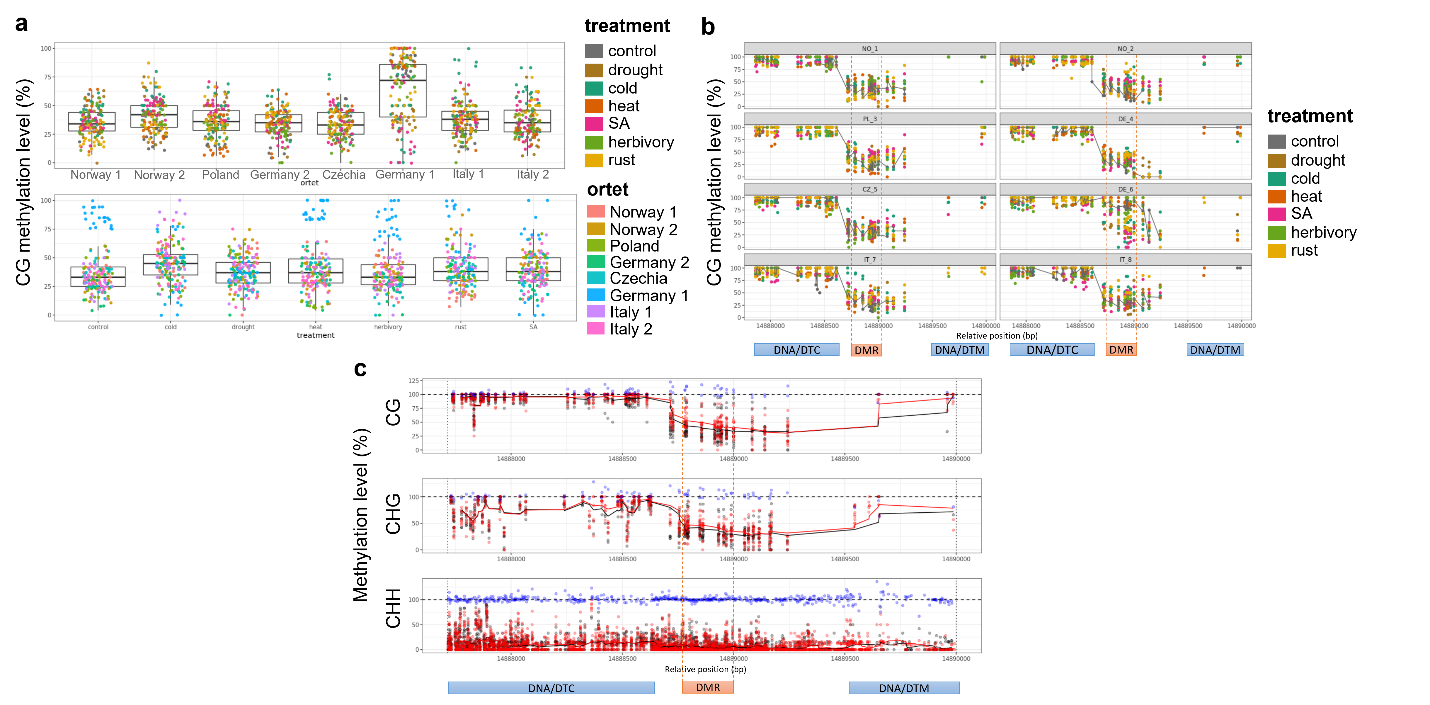

**Fig. S18** Methylation analysis of a hyper CG-DMR induced by cold and SA treatment. a) Boxplots of the methylation status of all CGs in the DMR, each dot represents a single CG and its methylation status within the DMR. Upper and bottom panels show the variation within ortet and treatment, respectively. b) Each panel shows the methylation profile of the DMR and DMR flanking regions for the respective ortet. Each dot represents a single CG, its methylation status and treatment. c) Each panel shows the methylation profile for each sequence context of the DMR and DMR flanking regions. Each dot represents a single cytosine, its methylation status and treatment (red = cold, black = control). Simple moving averages (SMA) over a period of 5 cytosines are also plotted (red line = cold, black line = control). Blue dots represent the difference between cold and control at each given position but using 100 as base line for better visualization. *The genomic context is shown below the panels: blue and red boxes represent transposable elements and DMR, respectively. Additionally, vertical dotted lines indicate the DMR edges.

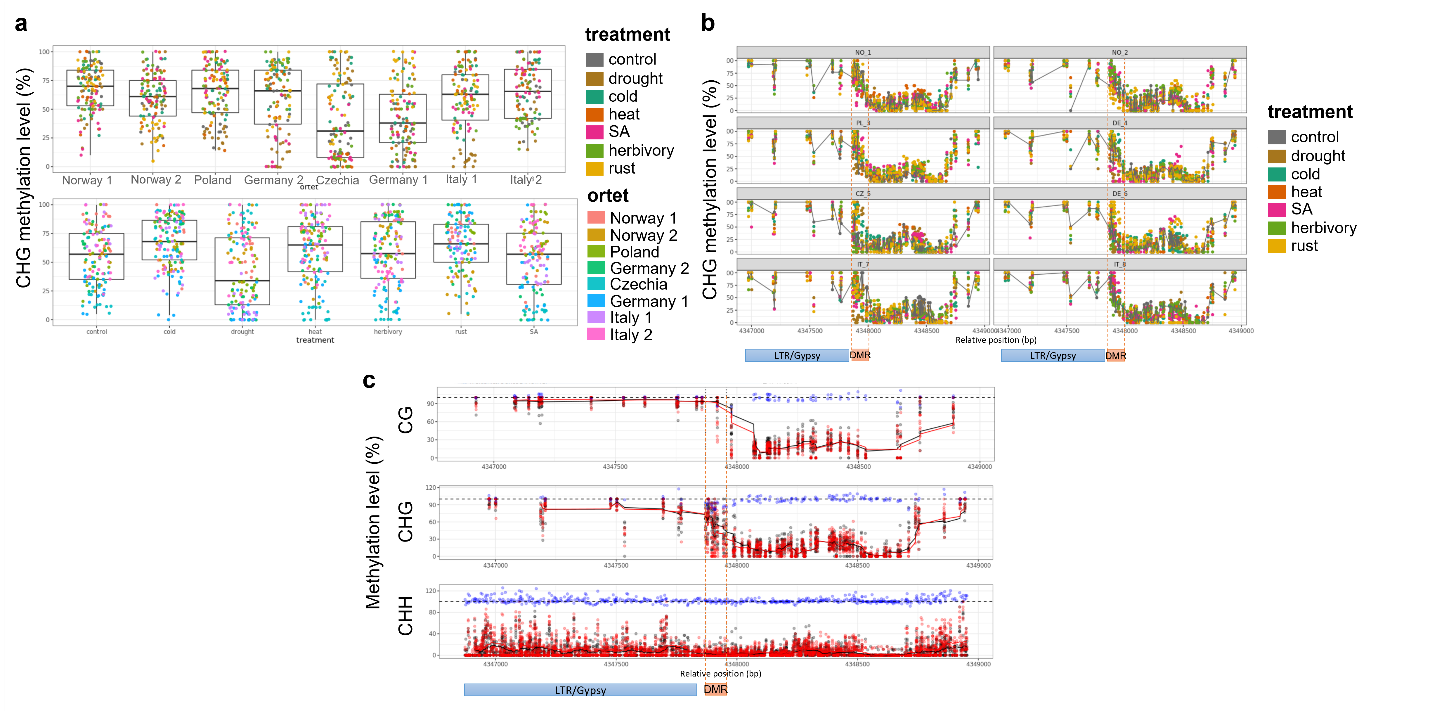

**Fig. S19** Methylation analysis of a hypo CHG-DMR induced by drought treatment. a) Boxplots of the methylation status of all CHGs in the DMR, each dot represents a single CHG and its methylation status within the DMR. Upper and bottom panels show the variation within ortet and treatment, respectively. b) Each panel shows the methylation profile of the DMR and DMR flanking regions for the respective ortet. Each dot represents a single CHG, its methylation status and treatment. c) Each panel shows the methylation profile for each sequence context of the DMR and DMR flanking regions. Each dot represents a single cytosine, its methylation status and treatment (red = drought, black = control). Simple moving averages (SMA) over a period of 5 cytosines are also plotted (red line = drought, black line = control). Blue dots represent the difference between drought and control at each given position but using 100 as base line for better visualization. *The genomic context is shown below the panels: blue and red boxes represent transposable elements and DMR, respectively. Additionally, vertical dotted lines indicate the DMR edges.

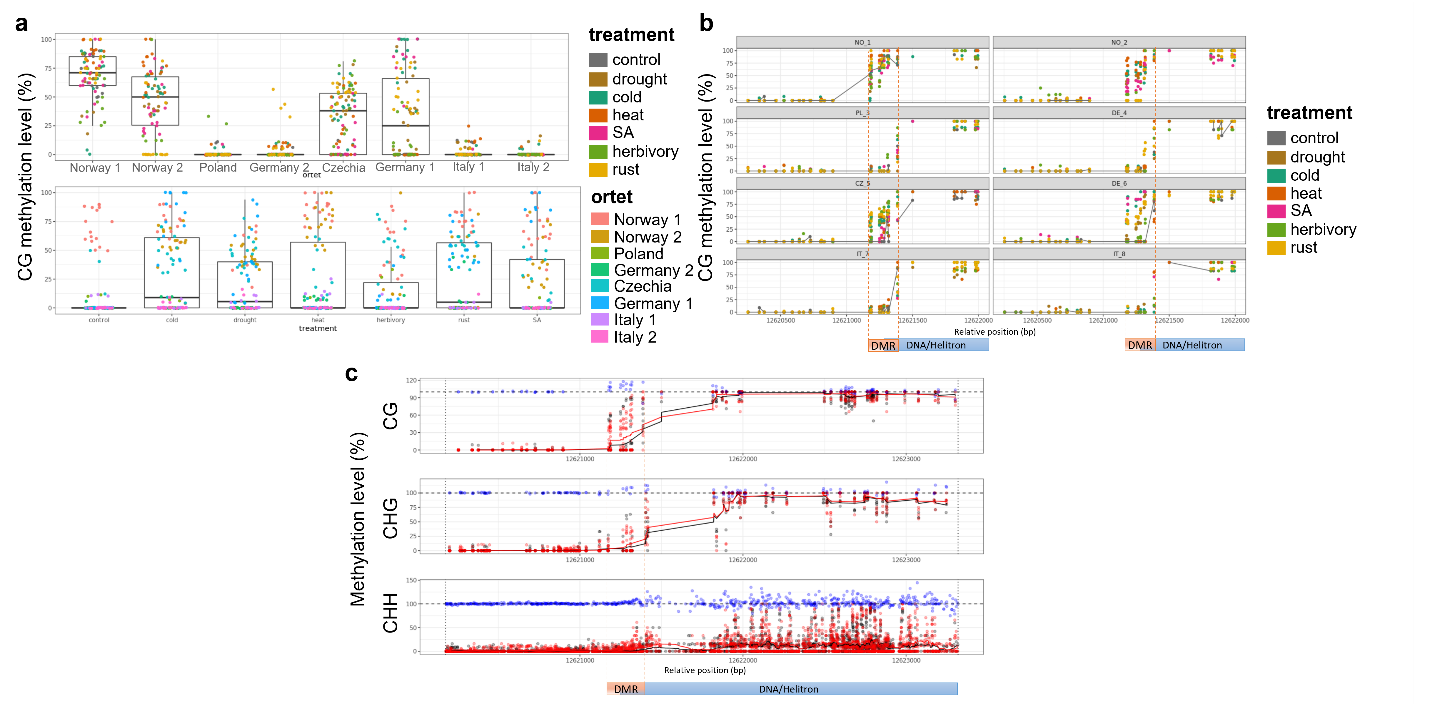

**Fig. S20**  Methylation analysis of a hyper CG-DMR induced by cold, drought and rust infection treatment. a) Boxplots of the methylation status of CGs in the DMR, each dot represents a single CG and its methylation status within the DMR. Upper and bottom panels show the variation within ortet and treatment, respectively. b) Each panel shows the methylation profile of the DMR and DMR flanking regions for the respective ortet. Each dot represents a single CG, its methylation status and treatment. c) Each panel shows the methylation profile for each sequence context of the DMR and DMR flanking regions. Each dot represents a single cytosine, its methylation status and treatment (red = cold, black = control). Simple moving averages (SMA) over a period of 5 cytosines are also plotted (red line = cold, black line = control). Blue dots represent the difference between cold and control at each given position but using 100 as base line for better visualization. *The genomic context is shown below the panels: blue and red boxes represent transposable elements and DMR, respectively. Additionally, vertical dotted lines indicate the DMR edges.

**Table S1** Description and geolocation of the ortets from which ramets were collected.

| **Ortet ID** | **Latitude** | **Longitude** | **European region** | **Country** | **Site** |
| --- | --- | --- | --- | --- | --- |
| **Norway 1** | 59.91953 | 10.77228 | North | Norway | Oslo Botanical Garden |
| **Norway 2** | 59.20995 | 10.94738 | North | Norway | Fredrikstad |
| **Poland** | 52.72787 | 19.11867 | Central east | Poland | Fabianci |
| **Czechia** | 50.08719 | 14.31551 | Central east | Czechia | Prague West |
| **Germany 1** | 49.50706 | 8.30128 | Central | Germany | Lambsheim |
| **Germany 2** | 52.70788 | 13.13274 | Central | Germany | Marwitz Ost |
| **Italy 1** | 44.60751 | 10.97689 | South | Italy | San Damaso |
| **Italy 2** | 44.56590 | 11.19290 | South | Italy | Budrie |

**Table S2** Data filtering and resolution of the methylation analyses. Sequencing data was filtered before using it as input for each analysis. Depending on the analysis, different resolution was targeted.

|  | **Genome-wide methylation** | | | | | **Differential methylation** | |
| --- | --- | --- | --- | --- | --- | --- | --- |
|  | **Average global methylation** | **PCA, HC** | **ICC** | **Methylation profiles (genes, TEs)** | **Methylation profiles (scaffold 1)** | **stress-DMR** | **ortet-DMR** |
| **Samples included in the analysis** | 52 | 52 | 52 | 56 | 56 | 56 | 56 |
| **low-coverage cytosines (removed)** | ≤ 5 | ≤ 5 | ≤ 5 | ≤ 5 | ≤ 5 | ≤ 5 | ≤ 5 |
| **low-coverage samples (removed)** | 4 | 4 | 4 | 0 | 0 | 0 | 0 |
| **Resolution** | Single base | Single base | 100-bp bins | Single base | 50-kb bins | Single base | Single base |

**Table S3** Summary of number of significant DMRs identified with the jack-knife approach (JK), Methylkit (M), and the intersection of both datasets

|  |  | **CpG** | | | |  | **CHG** | | | |  | **CHH** | | | |
| --- | --- | --- | --- | --- | --- | --- | --- | --- | --- | --- | --- | --- | --- | --- | --- |
| **TREATMENT** |  | **JK** | **intersection JK-M** | **M** | **JK verified with M (%)** |  | **JK** | **intersection JK-M** | **M** | **JK verified with M (%)** |  | **JK** | **intersection JK-M** | **M** | **JK verified with M (%)** |
| cold |  | 63 | 56 | 3078 | 88.89 |  | 81 | 77 | 4682 | 95.06 |  | 21 | 16 | 3766 | 76.19 |
| drought |  | 76 | 65 | 3596 | 85.53 |  | 140 | 128 | 7110 | 91.43 |  | 861 | 823 | 96777 | 95.59 |
| heat |  | 57 | 45 | 2304 | 78.95 |  | 79 | 68 | 4365 | 86.08 |  | 116 | 108 | 7698 | 93.10 |
| herbivory |  | 32 | 24 | 2108 | 75.00 |  | 21 | 18 | 2611 | 85.71 |  | 2 | 2 | 4060 | 100.00 |
| rust |  | 43 | 31 | 3151 | 72.09 |  | 77 | 69 | 3817 | 89.61 |  | 3 | 2 | 2804 | 66.67 |
| SA |  | 45 | 29 | 3246 | 64.44 |  | 79 | 69 | 5263 | 87.34 |  | 2 | 0 | 2183 | 0.00 |
| average |  |  |  |  | 77.48 |  |  |  |  | 89.21 |  |  |  |  | 71.92 |

**Table S4** Summary of significant DMRs classified according to the methylation direction compared to control group. For each context, percentages (%) refer to the relative amount of DMRs induced by each treatment. Ratios (hypo/hyper) represent the relative amount of hypomethylated DMRs compared to the number of hypermethylated DMRs.

|  | **CpG** | | | |  | **CHG** | | | |  | **CHH** | | | |  | **TOTAL** | |
| --- | --- | --- | --- | --- | --- | --- | --- | --- | --- | --- | --- | --- | --- | --- | --- | --- | --- |
|  | **HYPO** | **HYPER** | **%** | **hypo/hyper** |  | **HYPO** | **HYPER** | **%** | **hyper/hypo** |  | **HYPO** | **HYPER** | **%** | **hyper/hypo** |  | **HYPER** | **HYPO** |
| Cold | 9 | 54 | 19.94 | 0.17 |  | 3 | 78 | 16.98 | 0.04 |  | 10 | 11 | 2.09 | 0.91 |  | 143 | 22 |
| Drought | 58 | 18 | 24.05 | 3.22 |  | 121 | 19 | 29.35 | 6.37 |  | 0 | 861 | 85.67 | 0.00 |  | 898 | 179 |
| Heat | 35 | 22 | 18.04 | 1.59 |  | 49 | 30 | 16.56 | 1.63 |  | 10 | 106 | 11.54 | 0.09 |  | 158 | 94 |
| Herbivory | 16 | 16 | 10.13 | 1.00 |  | 11 | 10 | 4.40 | 1.10 |  | 1 | 1 | 0.20 | 1.00 |  | 27 | 28 |
| Rust | 21 | 22 | 13.61 | 0.95 |  | 48 | 29 | 16.14 | 1.66 |  | 2 | 1 | 0.30 | 2.00 |  | 52 | 71 |
| SA | 28 | 17 | 14.24 | 1.65 |  | 65 | 14 | 16.56 | 4.64 |  | 2 | 0 | 0.20 | - |  | 31 | 95 |
| SUM | 167 | 149 |  | 1.12 |  | 297 | 180 |  | 1.65 |  | 25 | 980 |  | 0.03 |  | 1309 | 489 |
| TOTAL | 316 | | 100 |  |  | 477 | | 100 |  |  | 1005 | | 100 |  |  | 1798 | |

**Table S5** Summary of significant DMRs classified according to stress specificity. For each context, percentages (%) refer to the relative amount of DMRs induced by single treatments (stress-specific) and several treatments (multi-stress). Ratios (M/S=Multi-stress/Stress-specific) represent the relative amount of multi-stress DMRs compared to the number of stress-specific DMRs.

|  | **CpG** | | |  | **CHG** | | |  | **CHH** | | |
| --- | --- | --- | --- | --- | --- | --- | --- | --- | --- | --- | --- |
|  | **Specific** | **Multi-stress** | **M/S ratio** |  | **Specific** | **Multi-stress** | **M/S ratio** |  | **Specific** | **Multi-stress** | **M/S ratio** |
| **Cold** | 32 (14.16%) | 31 | 0.97 |  | 43 (14.68%) | 38 | 0.88 |  | 10 (1.07%) | 11 | 1.10 |
| **Drought** | 36 (15.93%) | 40 | 1.11 |  | 73 (24.91%) | 67 | 0.92 |  | 804 (86.08%) | 57 | 0.07 |
| **Heat** | 23 (10.18%) | 34 | 1.48 |  | 34 (11.60%) | 45 | 1.32 |  | 61 (6.53%) | 55 | 0.90 |
| **Herbivory** | 13 (5.75%) | 19 | 1.46 |  | 6 (2.05%) | 15 | 2.50 |  | 2 (0.21%) | 0 | 0.00 |
| **Rust** | 17 (7.52%) | 26 | 1.53 |  | 37 (12.63%) | 40 | 1.08 |  | 2 (0.21%) | 1 | 0.50 |
| **SA** | 21 (9.29%) | 24 | 1.14 |  | 34 (11.60%) | 45 | 1.32 |  | 2 (0.21%) | 0 | 0.00 |
| **SUM** | 142 | 84 (37.17%) |  |  | 227 | 66 (22.53%) |  |  | 881 | 53 (5.67%) |  |
| **TOTAL** | 226 (100%) | |  |  | 293 (100%) | |  |  | 934 (100%) | |  |

**Table S6**  Z-test for proportion of DMRs on different genomic regions (Ha: p1 > p2). DMRs from all stress treatments were merged per context

| **CpG** | **#DMR** | **n1** | **p1** | **bp** | **n2 (total bp)** | **p2** | **pooled p** | **z statistic** | **p-value** |
| --- | --- | --- | --- | --- | --- | --- | --- | --- | --- |
| upstream 2kb | 26 | 226 | 0.1150 | 72912648 | 417170123 | 0.1748 | 0.1748 | -2.3646 | 0.9910 |
| gene body | 91 | 226 | 0.4027 | 137945985 | 417170123 | 0.3307 | 0.3307 | 2.3002 | **0.0107*** |
| downstream 2kb | 35 | 226 | 0.1549 | 70254286 | 417170123 | 0.1684 | 0.1684 | -0.5439 | 0.7067 |
| Introns | 34 | 226 | 0.1504 | 79918939 | 417170123 | 0.1916 | 0.1916 | -1.5712 | 0.9419 |
| Exons | 73 | 226 | 0.3230 | 58082296 | 417170123 | 0.1392 | 0.1392 | 7.9807 | **<0.001*** |
| TE | 142 | 226 | 0.6283 | 136755244 | 417170123 | 0.3278 | 0.3278 | 9.6237 | **<0.001*** |
| Intergenic | 85 | 226 | 0.3761 | 178172862 | 417170123 | 0.4271 | 0.4271 | -1.5497 | 0.9394 |
| Genic | 141 | 226 | 0.6239 | 238997261 | 417170123 | 0.5729 | 0.5729 | 1.5497 | 0.0606 |
| **CHG** | **#DMR** | **n1** | **p1** | **bp** | **n2 (total bp)** | **p2** | **pooled p** | **z statistic** | **p-value** |
| upstream 2kb | 21 | 267 | 0.0787 | 72912648 | 417170123 | 0.1748 | 0.1748 | -4.1359 | 1.0000 |
| gene body | 74 | 267 | 0.2772 | 137945985 | 417170123 | 0.3307 | 0.3307 | -1.8588 | 0.9685 |
| downstream 2kb | 32 | 267 | 0.1199 | 70254286 | 417170123 | 0.1684 | 0.1684 | -2.1202 | 0.9830 |
| Introns | 43 | 267 | 0.1610 | 79918939 | 417170123 | 0.1916 | 0.1916 | -1.2674 | 0.8975 |
| Exons | 45 | 267 | 0.1685 | 58082296 | 417170123 | 0.1392 | 0.1392 | 1.3834 | 0.0833 |
| TE | 198 | 267 | 0.7416 | 136755244 | 417170123 | 0.3278 | 0.3278 | 14.4026 | **<0.001*** |
| Intergenic | 147 | 267 | 0.5506 | 178172862 | 417170123 | 0.4271 | 0.4271 | 4.0784 | **<0.001*** |
| Genic | 120 | 267 | 0.4494 | 238997261 | 417170123 | 0.5729 | 0.5729 | -4.0784 | 1.0000 |
| **CHH** | **#DMR** | **n1** | **p1** | **bp** | **n2 (total bp)** | **p2** | **pooled p** | **z statistic** | **p-value** |
| upstream 2kb | 211 | 932 | 0.2264 | 72912648 | 417170123 | 0.1748 | 0.1748 | 4.1491 | **<0.001*** |
| gene body | 162 | 932 | 0.1738 | 137945985 | 417170123 | 0.3307 | 0.3307 | -10.1783 | 1.0000 |
| downstream 2kb | 188 | 932 | 0.2017 | 70254286 | 417170123 | 0.1684 | 0.1684 | 2.7174 | **0.0033*** |
| Introns | 127 | 932 | 0.1363 | 79918939 | 417170123 | 0.1916 | 0.1916 | -4.2905 | 1.0000 |
| Exons | 41 | 932 | 0.0440 | 58082296 | 417170123 | 0.1392 | 0.1392 | -8.3986 | 1.0000 |
| TE | 853 | 932 | 0.9152 | 136755244 | 417170123 | 0.3278 | 0.3278 | 38.2029 | **<0.001*** |
| Intergenic | 377 | 932 | 0.4045 | 178172862 | 417170123 | 0.4271 | 0.4271 | -1.3943 | 0.9184 |
| Genic | 555 | 932 | 0.5955 | 238997261 | 417170123 | 0.5729 | 0.5729 | 1.3943 | 0.0816 |

**Table S7**  Contingency tables of DMRs according to TE and genomic feature associations. Genic region stands for the gene body plus 2 kb upstream and 2kb downstream regions. Each sequence context was tested separately. First, Chi-square tests for independence were performed, then after significant differences were obtained, differences between the marginal proportions (McNemar’s test) were also evaluated

|  | **CpG** | |  | **CHG** | |  | **CHH** | |
| --- | --- | --- | --- | --- | --- | --- | --- | --- |
|  | **no TE** | **TE** |  | **no TE** | **TE** |  | **no TE** | **TE** |
| **intergenic** | 11 | 74 |  | 20 | 127 |  | 26 | 351 |
| **genic** | 73 | 68 |  | 49 | 71 |  | 53 | 502 |
| Chi-Sqr test | <0.001* | |  | **<0.001*** | |  | 0.564 | |
| McNemar's test | 1 | |  | **<0.001*** | |  | <0.001* | |
|  | **no TE** | **TE** |  | **no TE** | **TE** |  | **no TE** | **TE** |
| **intron** | 17 | 17 |  | 17 | 26 |  | 10 | 117 |
| **exon** | 46 | 27 |  | 29 | 16 |  | 18 | 23 |
| Chi-Sqr test | 0.654 | |  | 0.141 | |  | **<0.001*** | |
| McNemar's test | <0.001* | |  | 0.787 | |  | **<0.001*** | |
|  | **no TE** | **TE** |  | **no TE** | **TE** |  | **no TE** | **TE** |
| **upstream 2k** | 10 | 14 |  | 6 | 13 |  | 16 | 194 |
| **gene body** | 55 | 36 |  | 37 | 37 |  | 25 | 137 |
| Chi-Sqr test | 0.436 | |  | 0.559 | |  | 0.127 | |
| McNemar's test | <0.001* | |  | 0.001* | |  | **<0.001*** | |
|  | **no TE** | **TE** |  | **no TE** | **TE** |  | **no TE** | **TE** |
| **downstream 2k** | 8 | 18 |  | 6 | 21 |  | 12 | 171 |
| **gene body** | 55 | 36 |  | 37 | 37 |  | 25 | 137 |
| Chi-Sqr test | 0.067 | |  | 0.100 | |  | 0.069 | |
| McNemar's test | <0.001* | |  | 0.048* | |  | **<0.001*** | |

**Table S8** Fold enrichment analysis of TE superfamilies targeted by drought-induced CHH DMRs. The total TE length of 136’755.244 bp was used to calculate TE proportions. P-values were adjusted using Bonferroni correction based on the number of tested superfamilies.

| **TE superfamily** | **TE length (bp)** | **TE proportion** | **#DMR** | **DMR proportion** | **fold enrichment** | **log2 fold enrichment** | **hypergeometric test (P-value)** | **Adjusted**  **P-value** |
| --- | --- | --- | --- | --- | --- | --- | --- | --- |
| DNA/DTA | 4160939 | 0.0304 | 29 | 0.0383 | 1.2574 | 0.3305 | 0.127 | 1.000 |
| DNA/DTC | 12503807 | 0.0914 | 50 | 0.0660 | 0.7214 | -0.4710 | 0.995 | 1.000 |
| DNA/DTH | 3166333 | 0.0232 | 27 | 0.0356 | 1.5384 | 0.6215 | 0.020 | 0.324 |
| DNA/DTM | 16517837 | 0.1208 | 57 | 0.0752 | 0.6226 | -0.6837 | 1.000 | 1.000 |
| DNA/DTT | 949090 | 0.0069 | 10 | 0.0132 | 1.9009 | 0.9267 | 0.042 | 0.667 |
| DNA/Helitron | 41685589 | 0.3048 | 135 | 0.1781 | 0.5843 | -0.7753 | 1.000 | 1.000 |
| LTR/Copia | 11836722 | 0.0866 | 62 | 0.0818 | 0.9450 | -0.0816 | 0.698 | 1.000 |
| LTR/Gypsy | 37558006 | 0.2746 | 119 | 0.1570 | 0.5716 | -0.8068 | 1.000 | 1.000 |
| LTR/unknown | 12388115 | 0.0906 | 69 | 0.0910 | 1.0049 | 0.0070 | 0.501 | 1.000 |
| MITE/DTA | 668757 | 0.0049 | 2 | 0.0026 | 0.5396 | -0.8902 | 0.885 | 1.000 |
| MITE/DTC | 172232 | 0.0013 | 5 | 0.0066 | 5.2376 | 2.3889 | 0.003 | **0.048*** |
| MITE/DTH | 572132 | 0.0042 | 27 | 0.0356 | 8.5142 | 3.0899 | <0.001 | **<0.001*** |
| MITE/DTM | 478225 | 0.0035 | 3 | 0.0040 | 1.1318 | 0.1786 | 0.495 | 1.000 |
| MITE/DTT | 61212 | 0.0004 | 0 | 0.0000 | 0.0000 | NULL | 1.000 | 1.000 |
| SINE | 471999 | 0.0035 | 148 | 0.1953 | 56.5712 | 5.8220 | <0.001 | **<0.001*** |
| target_site_duplication | 2011758 | 0.0147 | 15 | 0.0198 | 1.3452 | 0.4278 | 0.156 | 1.000 |

**Table S9** Summary of DMRs identified between each pair of ortets on each sequence context. Color gradient highlights pairs with the largest differences.

| **CpG** | **NO_1** | **NO_2** | **PL_3** | **DE_4** | **CZ_5** | **DE_6** | **IT_7** | **IT_8** | **SUM** |
| --- | --- | --- | --- | --- | --- | --- | --- | --- | --- |
| **NO_1** |  | 1548 | 902 | 818 | 1274 | 1762 | 1529 | 1358 | 9191 |
| **NO_2** | 1548 |  | 1397 | 1497 | 1677 | 1788 | 2100 | 1717 | 11724 |
| **PL_3** | 902 | 1397 |  | 794 | 1129 | 1656 | 1385 | 1226 | 8489 |
| **DE_4** | 818 | 1497 | 794 |  | 1119 | 1702 | 1365 | 1341 | 8636 |
| **CZ_5** | 1274 | 1677 | 1129 | 1119 |  | 1169 | 1406 | 1302 | 9076 |
| **DE_6** | 1762 | 1788 | 1656 | 1702 | 1169 |  | 1911 | 1666 | 11654 |
| **IT_7** | 1529 | 2100 | 1385 | 1365 | 1406 | 1911 |  | 1380 | 11076 |
| **IT_8** | 1358 | 1717 | 1226 | 1341 | 1302 | 1666 | 1380 |  | 9990 |
|  | **avg #DMRs per comparison** | | | | | | | 1425 |  |
| **CHG** | **NO_1** | **NO_2** | **PL_3** | **DE_4** | **CZ_5** | **DE_6** | **IT_7** | **IT_8** | **SUM** |
| **NO_1** |  | 2017 | 1223 | 946 | 1782 | 2129 | 1305 | 1266 | 10668 |
| **NO_2** | 2017 |  | 2007 | 1944 | 2318 | 2014 | 2483 | 1890 | 14673 |
| **PL_3** | 1223 | 2007 |  | 1078 | 1598 | 1922 | 1366 | 1160 | 10354 |
| **DE_4** | 946 | 1944 | 1078 |  | 1242 | 1849 | 1060 | 1212 | 9331 |
| **CZ_5** | 1782 | 2318 | 1598 | 1242 |  | 1399 | 1367 | 1640 | 11346 |
| **DE_6** | 2129 | 2014 | 1922 | 1849 | 1399 |  | 1983 | 1833 | 13129 |
| **IT_7** | 1305 | 2483 | 1366 | 1060 | 1367 | 1983 |  | 1368 | 10932 |
| **IT_8** | 1266 | 1890 | 1160 | 1212 | 1640 | 1833 | 1368 |  | 10369 |
|  | **avg #DMRs per comparison** | | | | | | | 1621 |  |
| **CHH** | **NO_1** | **NO_2** | **PL_3** | **DE_4** | **CZ_5** | **DE_6** | **IT_7** | **IT_8** | **SUM** |
| **NO_1** |  | 226 | 51 | 54 | 69 | 143 | 78 | 58 | 679 |
| **NO_2** | 226 |  | 272 | 248 | 191 | 110 | 350 | 270 | 1667 |
| **PL_3** | 51 | 272 |  | 35 | 80 | 254 | 132 | 61 | 885 |
| **DE_4** | 54 | 248 | 35 |  | 37 | 187 | 103 | 76 | 740 |
| **CZ_5** | 69 | 191 | 80 | 37 |  | 66 | 92 | 74 | 609 |
| **DE_6** | 143 | 110 | 254 | 187 | 66 |  | 179 | 184 | 1123 |
| **IT_7** | 78 | 350 | 132 | 103 | 92 | 179 |  | 60 | 994 |
| **IT_8** | 58 | 270 | 61 | 76 | 74 | 184 | 60 |  | 783 |
|  | **avg #DMRs per comparison** | | | | | | | 133 |  |

**Table S10** Summary of ortet-DMRs and the intersection with stress-DMRs. DMRs were further classified based on its uniqueness, i.e., the frequency they appeared on individual pairwise comparisons: identified on a unique comparison or in more than one comparison.

|  |  |  |  | **stress-DMRs** | | | |
| --- | --- | --- | --- | --- | --- | --- | --- |
| **CpG** | **ortet-DMRs** | **%** |  | **with ortet-DMR intersection** | **%** |  | **no ortet-DMR intersection** |
| unique comparison | 3044 | 30.93 |  | 21 | 13.04 |  |  |
| more than one comparison | 6796 | 69.07 |  | 140 | 86.96 |  |  |
| TOTAL | 9840 |  |  | 161 (71.24%) |  |  | 65 (28.76%) |
|  |  |  |  | 226 (100%) | | | |
| **CHG** | **ortet-DMRs** | **%** |  | **with ortet-DMR intersection** | **%** |  | **no ortet-DMR intersection** |
| unique comparison | 1529 | 20.79 |  | 32 | 11.03 |  |  |
| more than one comparison | 5824 | 79.21 |  | 258 | 88.97 |  |  |
| TOTAL | 7353 |  |  | 290 (85.29%) |  |  | 50 (14.71%) |
|  |  |  |  | 340 (100%) | | | |
| **CHH** | **ortet-DMRs** | **%** |  | **with ortet-DMR intersection** | **%** |  | **no ortet-DMR intersection** |
| unique comparison | 501 | 43.91 |  | 22 | 36.67 |  |  |
| more than one comparison | 640 | 56.09 |  | 38 | 63.33 |  |  |
| TOTAL | 1141 |  |  | 60 (6.36%) |  |  | 884 (93.64%) |
|  |  |  |  | 944 (100%) | | | |

**Methods S1** **Plant material.** Between March and May 2018, cuttings from eight adult *Populus nigra* cv. ‘Italica’ clones were collected from five European countries (Table S1, Figure 1A; see Díez-Rodríguez et al., 2022). At each site, at least seven hardwood cuttings (ramets) of approximately 30 cm length and 20 mm diameter were sampled from each parental tree (ortet) and stored at 4 °C for two weeks until planting. In order to promote root development, cuttings were sliced at the base (1 cm) before soaking in water to ¾ of their length for three days in darkness (with the axillary bud scales upright). Then, cuttings were grown under controlled greenhouse conditions for 12 weeks until the start of the experiment. Growth conditions were: 22/18 °C (±2°C) at day/night, 60% Relative humidity (±5% Rh), 16/8 h light/dark enabled by additional lighting (when needed) with high pressure sodium lamps Son-T, 600W Philips GP to provide a minimum of 250 µmol/m^2^/s PAR. Cuttings were planted first in 4-liter pots with 3:1 sand:peat mixture (v/v). The sand consisted of two parts of coarse sand (0.7-1.25 mm grain size), and one part of fine sand (0.4-0.8 mm), while peat referred to nutrient-poor potting soil (“zaai- en stek, Lensli substrates”). Pots were placed in a flood table for three weeks. Rooted ramets of similar size (shoot length > 15 cm) were transferred to 7-liter pots with 1:1 sand:peat mixture (30% coarse sand, 20% fine sand, and 50% nutrient-poor potting soil) and maintained with regular watering to pot capacity (3-5 times per week). Two weeks prior to the start of the experiment, three grams of slow-release fertilizer Osmocote Exact Mini (16+8+11+2MgO+TE) were added to each pot.

**Methods S2** **Experimental design.** Seven 3-month-old ramets of similar size were selected from each of the eight (ortets) for subsequent exposure to different environmental treatments. The experiment consisted of three biotic and three abiotic stresses plus a single control group, with eight replicate plants per treatment (56 trees in total), where each parental tree contributed one replicate to each of the treatments. All treatments were implemented simultaneously during a period of 25 days, including two 10-day stress periods and a 5-day stress-free period in-between (Figure S1). **Control group:** during the entire stress experiment control plants were maintained in a greenhouse with controlled conditions as stated above. Plants were watered daily to pot capacity; thus, soil volumetric water content (VWC) was maintained on average at 20.2% (± 3.24 SD). VWC was monitored daily using WET Sensor kit (Delta-T Devices) which includes WET Sensor and HH2 Moisture Meter. Two measurements were performed per pot and the mean VWC was calculated for each plant (Figure S2). During the entire experiment control plants were maintained in the same greenhouse table with other stress treatments in a Latin square design unless otherwise stated (Figure S3). **Rust infection:** Uredospores of the poplar leaf rust fungus (*Melampsora larici-populina* Kleb.) were obtained from Dr. Sybille Unsicker (Max Planck Institute for Chemical Ecology, Jena, Germany). Spore collection, amplification and identification are described in Eberl et al., (2018). Spores were stored at -20°C until the start of the experiment. On experimental day 0, one branch of each plant with 15-20 leaves was completely covered with a polyethylene terephthalate bag (Bratschlauch, Toppits, Minden, Germany). Then, enclosed leaves were spray-inoculated with 3 ml of spore-water solution (1 mg/ml) on the abaxial side. On day 11, the entire infected branch was cut off. On day 15, in the same manner a second rust inoculation was applied on approximately 20 leaves (from the fifth mature leaf downwards) of the main branch by spray-inoculation. In both exposure periods, the bags remained closed to ensure sufficient humidity and to avoid cross contamination, however small holes were made in the bag to enable some aeration. First sporangia in the rust-infected leaves were visible on the abaxial side of the leaves at 7 dpi (days post-infection). No signs of infected leaves outside the bags were observed during the experiment. Plants were maintained in a separate greenhouse with the same environmental and watering conditions as the control group to avoid direct contact with control plants. **Herbivory treatment:** Gypsy moth (*Lymantria dispar* L.) caterpillars (L2) were obtained from Dr. Sybille Unsicker (Max Planck Institute for Chemical Ecology, Jena, Germany), and maintained in a climate chamber at 14/10 h light/dark, 20°C, and 60% humidity until the start of the experiment. Caterpillars were fed first with artificial diet (MP Biomedicals LLC) and then with poplar leaves for two days prior to the start of the experiment. On experimental day 1, the smaller branch (15-20 leaves) of each plant was completely enclosed in an anti-insect mesh bag. Then, eight L3 caterpillars were carefully placed inside the bag. On day 11, both caterpillars and mesh bags were removed. On day 16, approximately 20 leaves (from the fifth mature leaf downwards) on the main branch were enclosed in an anti-insect mesh bag, then ten L3-L4 caterpillars were placed inside the bag. No signs of eaten leaves outside the bags were observed during the experiment. Plants were maintained in a separate greenhouse with the same environmental and watering conditions as the control group to avoid direct contact with control plants. **Salicylic acid (SA) treatment:**  During the two stress periods from days 1-10, and 16-25, once a day, of each plant three random leaves of different size were sprayed on both sides with a total of 2 ml of 1 mM SA (Sigma-Aldrich) containing 0.1% Triton X-100 (Sigma-Aldrich). The ten youngest leaves of the main branch were not used for SA treatment. Plants were maintained in a separate greenhouse with the same environmental and watering conditions as the control group to avoid direct contact with control plants.  **Drought stress:** VWC was monitored daily using WET Sensor kit (Delta-T Devices). Two measurements were performed per pot and the mean VWC was calculated. Plants were gradually dried by withholding watering for two days until a VWC of 8% was reached at experimental day 1. VWC was maintained at an average of 8.19% (± 2.77 SD) during the stress periods (days: 1-10, 16-25). Plants, whose VWC dropped below 6% were watered to reach the mean VWC of 8%. During the stress-free period (days: 11-15), VWC was raised to 17.8% (± 2.27 SD) by daily watering to pot capacity (Figure S2). During the entire experiment plants were maintained on the same greenhouse table as the control group in a Latin square design (Figure S3). **Heat stress:** During both stress periods (days: 1-10, 16-25), plants were transferred to a climate chamber with humidity 60%, 16/8 h light/dark (high pressure Sodium lamps Son-T, 600W Philips GP, 350 µmol/m2/s PAR) and temperature was adjusted as follows: 30/28°C (day/night) for 4 days, 38/28°C for 3 days, and 30/28°C for 3 days. Due to the high evaporation rates, plants were watered daily to maintain VWC close to control conditions (on average 16.66%, ± 6.58 SD, based on daily measurements as described above). During the stress-free period, plants were moved to the same greenhouse table as the control group in a Latin square design (Figure S3).  **Cold stress:** During the periods of stress exposure (days: 1-10, 16-25), plants to be treated by cold stress were moved to a climate chamber that was maintained at 4/4°C (day/night), humidity 60% and 16/8 h light/dark (high pressure Sodium lamps Son-T, 600W Philips GP, 50 µmol/m2/s PAR). VWC was maintained close to control conditions (20.8%, ± 2.51 SD) by regular watering to pot capacity. During the stress-free period, plants were moved to the same greenhouse table as the control group in a Latin square design (Figure S3).  **Harvesting:** For DNA methylation analysis, on experimental day 26, twelve circular punches (Ø 8 mm; ~ 100 mg fresh weight in total) were cut out from the eighth mature leaf (counting from the apex of the main branch, leaf plastochron index: 10) of each plant. Mid‐ribs were avoided, and leaf punches were immediately frozen in liquid nitrogen and stored at −80 °C. All leaves that were used for sample collection did not show signs of damage/infection at the time of sampling. Sampling order was determined by first randomizing ramets by region and then by treatment within each region. Branch length was measured at experimental day 1 and 29. Ten days after the end of the experiment all ramets were coppiced. Stems and leaves were dried at 70°C for 7 days, then dry weight biomass was determined. Plants were maintained with minimal watering (200 ml/day) from the end of the experiment until coppicing. The main stem diameter of each cutting was also measured after coppicing.

**Methods S3** **DNA extraction and whole genome bisulfite sequencing.** Per sample, frozen leaf tissue (~ 100 mg fresh weight per sample) was grinded and homogenized using TissueLyser II (QIAGEN), then genomic DNA was isolated using the Sodium Dodecyl Sulfate (SDS) procedure of the NucleoSpin Plant II DNA isolation kit (Macherey-Nagel, Dueren, Germany). DNA extraction order was determined by first randomizing samples by region and then by treatment within each region Preparation of DNA libraries for bisulfite sequencing was performed as described in (Nunn *et al.*, 2022). Briefly, 300 ng genomic DNA was fragmented to 350 bp average size with a Covaris S2 instrument. Then, libraries were prepared using the NEBNext Ultra II DNA Library Prep Kit for Illumina (New England Biolabs) with the following modifications. Adaptor dilution 1:2 before the adaptor ligation step (using NEBNext Methylated Adaptor E7535S for Illumina). AMPure XP Beads were used for size Selection (Approx. final library size: 480 bp). Non-methylated cytosine residues were converted to uracil using the EZ-96 DNA Methylation-Gold MagPrep (Zymo Research) according to the manufacturer’s guidelines. Library enrichment was performed with KAPA HiFi HotStart Uracil+ ReadyMix PCR Kit (Roche) and 12 PCR cycles. Library preparation order was determined by first randomizing samples by ortet and then by treatment within each ortet. All sequencing was performed by Novogene on an Illumina HiSeq X Ten sequencing system. Libraries were sequenced with 2x150-bp paired-end reads at 30X coverage. Libraries were sequenced in a total of eight sequencing lanes trying to allocate ramets derived from the same ortet in the same lane to avoid batch effects (sequencing summary in Supplementary file 1). The sodium bisulfite non-conversion rate was calculated as the percentage of cytosines sequenced at cytosine reference positions in the chloroplast genome.

**Methods S4** **Processing of bisulfite-treated reads and methylation calling**. Sequenced reads were trimmed, filtered, and aligned to the *de novo* reference genome using the EpiDiverse Toolkit (WGBS pipeline v1.0, <https://github.com/EpiDiverse/wgbs>) (Nunn et al., 2021). Briefly, low-quality ends from reads were trimmed (minimum base quality threshold: 20) using Cutadapt (<https://cutadapt.readthedocs.io/en/stable>), then sequencing adapters were recognized and removed (minimum overlap length for adapter sequences in reads: 3 bp). Trimmed reads shorter than 36 bases were discarded. The remaining high-quality reads (on average 99.85% of raw reads across all the samples) were aligned against the *Populus nig*r*a* var. Italica *de novo* reference genome (ENA project: PRJEB44889) using the bisulfite-treated sequences aligner erne-bs5 (<http://erne.sourceforge.net>) with the parameters as follows. Insert size between paired-end reads: 0-600 bp, proportion of allowed opposite-strand bisulfite mismatches: 0.05. Only reads mapping uniquely to a single position were used for this study (on average, 64% of filtered reads) (Supplementary file 1). Within the same EpiDiverse/WGBS pipeline, per-cytosine methylation metrics were extracted from the aligned reads using MethylDackel (<https://github.com/dpryan79/MethylDackel>). Three bedGraph files per sample were produced, corresponding to cytosines on each sequence context: CpG, CHG and CHH.

**Methods S5** **Principal component analysis, hierarchical clustering and correlation analysis.** For CpG and CHG context, the same filtered genomic positions used for average global methylation analysis were considered for unsupervised analyses: PCA and HC. For CHH context, an extra filtering step was conducted as most cytosines with methylation levels close to either 0% or 100% showed very low variation across samples. Only cytosines where at least 10% of the samples (5/52) showed methylation levels either higher than 5% or lower than 95% were retained for analysis (1’153.473 positions). Principal components were calculated in R using the *prcomp* function of the *stats* package. Hierarchical clustering (Ward’s method) was performed on the same data by first calculating the corresponding distance matrix (Manhattan method). R functions *dist* and *hclust* from *stats* package were used. Pairwise correlation analysis was performed using genomic regions rather than single positions. After removing outliers and low sequencing read depth positions as stated above, the poplar genome was compartmentalized in 100-bp non-overlapping bins. Each sequence context was analyzed separately. Bins containing four or less cytosines were removed from the analysis. Average methylation per bin was calculated, and only bins with methylation information across all (52) samples were retained. Finally, bins with very low methylation variation across samples (SD ≤ 1%) were removed. The intraclass correlation coefficient (ICC) reflects both degree of correlation and agreement between measurements (Koo & Li, 2016).  ICC was calculated for all pairwise comparisons between samples. The ICC takes a value from zero (implying no agreement) to 1 (perfect agreement). Coefficients were calculated in R using the *icc* function of the *irr* package. We used the ICC form: Two-way random effects, absolute agreement, single measurement, according to McGraw and Wong (1996) convention.

**Methods S6** **Methylation profiles.** All (56) samples were included in the analysis. For each sequence context, methylation distribution around genes and TEs was calculated using cytosines with at least six sequencing read depth. For calculation of methylation levels over gene regions, only protein-coding genes with known 5’UTR and 3’UTR coordinates were considered. For methylation over transposable elements, only TEs longer than 150 bp were analyzed in order to avoid noise from flanking regions of fragmented TEs, First, for each sample, methylation levels were extracted from: gene bodies, and 2 kb upstream/downstream flanking regions. Then, per-cytosine average methylation level was calculated across treatment replicates. Next, for each treatment, methylation levels at relative positions to the transcription start site (TSS) and transcription termination site (TTS) were averaged for all genes. Simple moving averages (SMA) over a period of 50 bp were calculated and plotted for each treatment and context. SMAs were calculated in R using the *geom_ma* function of the *tidyquant* package. The same procedure was used to analyze methylation levels over TE regions (TE bodies and flanking regions).

**Methods S7** **DMR calling.** As ramets had a strong “legacy” effect in their methylation background reflecting its ortet origin (see global methylation patterns in results section), having all eight ortets represented in each of the treatments contributes considerable within-group variation and hence reduces power for DMR identification. To minimize this effect, stress-DMRs were identified in a jack-knife approach (leave-one-out) as follows. For each treatment–control comparison, eight DMR callings were performed (Figure S4). On each DMR calling, one ortet was left out at a time by removing its corresponding treatment and control ramets. Thus, seven replicates (ortets) were included on each DMR calling; this procedure reduces the loss of statistical power that is caused by deviant methylation patterns in a single ortet. All identified DMRs were retained for further analysis, the same DMRs appearing in two or more DMR callings were merged as follows, start: left-most position, end: right-most position, control vs treatment difference: average, number of cytosines: average, DMR length: average. Additionally, in order to check if our jack-knife approach for DMR calling produced robust results, we performed another DMR calling using Methylkit (Akalin et al., 2012). Briefly, after methylation calling in all samples, cytosines with five or less sequencing read depth were removed. Then, the genome was split into 100-bp non-overlapped fragments. Genomic fragments containing less than six cytosines were removed from the analysis. For each treatment-control comparison, DMR calling was performed on fragments with at least six replicates per treatment (6/8), using ortet as covariate in the model. Significant DMRs (q-value < 0.05) found by Methylkit were filtered (minimum 5% difference) and then intersected with the set of DMRs called using jack-knife approach. Each sequence context was analyzed separately. This check indicated a large overlap between Methylkit-detected DMRs and the DMRs detected by our method, and also highlighted the conservative nature of our results (Table S2). Then, based on the genomic location, DMRs identified in more than one treatment were classified as multi-stress DMRs while the rest were labeled as stress-specific DMRs. Multi-context DMRs were also identified in the same manner. The methylation direction (hyper/hypo) of multi-stress DMRs was further examined since in certain regions some treatments induced opposite methylation responses. Regions with positive methylation difference with respect to control group were called hypermethylated DMRs while hypomethylated DMRs stood for negative methylation differences. We performed additional analyses to detect DMRs among ortets rather than among treatments. Ramets derived from the same ortet and exposed to different treatments were considered replicates. Briefly, DMRs were called for all pairwise comparisons among the eight ortets (total: 28 DMR sets per context). Next, for each context, we combined all DMRs using *bedtools* merge, and then, we classified each DMR according to the number of pairwise comparison where each DMR appeared (*unique comparison* or *shared by two or more comparisons*). Intersections between stress-DMRs and ortet-DMRs were performed using *bedtools* intersect.

**Methods S8** **DMR annotation.** Statistically significant DMRs (jack-knife approach) were annotated based on the *P. nigra* Italica protein-coding gene model annotation (ZENODO). First, DMRs were classified as genic (gene body, and within 2kb gene flanking regions) or intergenic DMRs (>2kb apart from genes) using *bedtools* intersect. Only the longest transcript per gene was used for this analysis.  For DMR distribution over genic regions, relative positions to TSS and TTS for DMRs located outside genes were calculated based on the distance from the DMR midpoint to the most proximal TSS or TTS (maximum 5kb). DMR locations within gene bodies were normalized to 2-kb gene length. Genic DMRs were further classified according to their location and overlaps with other gene features: “*UTR5p”* (upstream region and first exon), *“exon”,* “*exon-intron”*, “*intron”*, “*UTR3p”* (last exon and downstream region), and *“>2 features”* (upstream/downstream region, exon, and/or intron). For exon and intron intersections, overlaps of either 30% of DMR length or 70% of the genic feature were required. DMRs were also associated with transposable elements based mainly on the TE prediction available in ZENODO. Short interspersed nuclear elements (SINEs) were added to the predicted TEs based on BLASTN results (70% similarity, 90% coverage) using consensus sequences of Salicaceae SINE families (Kogle*r et al*. 2020). A DMR was associated with a transposable element when it was located inside or less than 200 bp away from a TE. Only the shortest and closest TE was retained when a DMR was associated with more than one TE. For DMR distribution over TE regions, relative positions to TE start site (TESS) and termination site (TETS) were calculated based on the distance from the DMR midpoint to the most proximal TESS or TETS (maximum 1.5 kb). DMR locations within TE bodies were normalized to 500-bp length. We combined all DMRs from all treatments (keeping only one DMR when two or more DMRs overlap) to test enrichment in genomic regions and to plot DMR distributions for each sequence context. Additionally, as most drought CHH-DMRs were associated to TEs, we calculated the fold enrichment for each TE superfamily considering the proportion of TE-associated DMRs on each TE superfamily compared to the proportion (in length) of each TE superfamily within the whole TE content in the black poplar reference genome. Only DMRs located inside TEs were analyzed. P-values were obtained from the hypergeometric test and then adjusted (Bonferroni) according to the number of TE superfamilies tested.

**Methods S9** **Gene ontology enrichment analysis.** Functional enrichment analysis has to be carefully interpreted as gene expression data was not collected in this experiment and most stresses produced very few DMRs. Therefore, our analysis was mainly focused on medium-large gene sets associated with drought-induced methylation responses.  Each DMR was associated with its overlapping gene and/or with the closest gene (maximum 2kb upstream from TSS). Genes associated to drought CHH-DMRs were subjected to gene ontology (GO) enrichment analysis. The gene background (universe) was built with the closest Arabidopsis (*A. thaliana*) homologue of each *P. nigra* Italica gene, which was determined using BLAST best reciprocal hits (RBH) of the protein sequences (R package *orthologr*). Best hits were filtered by keeping alignments covering at least 60% of both *Arabidopsis* and *P. nigra* proteins, and minimum 60% similarity. *Arabidopsis* sequence proteins were extracted from phytozome V13, and functional annotations were retrieved from the PLAZA 5.0 dicots database (<https://bioinformatics.psb.ugent.be/plaza/>). GO enrichments were performed using clusterProfiler v4 (Wu et al., 2021). P-values were adjusted for multiple testing controlling the positive false discovery rate (q-value).
